# *Akr1d1-/-* mice have a sexually dimorphic metabolic phenotype with reduced fat mass, increased insulin sensitivity and hypertriglyceridemia in males

**DOI:** 10.1101/2021.02.02.429227

**Authors:** Laura L Gathercole, Nikolaos Nikolaou, Anastasia Arvaniti, Shelley E Harris, Toryn M Poolman, Jonathan M Hazlehurst, Denise V Kratschmar, Marijana Todorčević, Ahmad Moolla, Niall Dempster, Ryan C Pink, Michael F Saikali, Liz Bentley, Trevor M Penning, Claes Ohlsson, Carolyn L Cummins, Matti Poutanen, Alex Odermatt, Roger D Cox, Jeremy W Tomlinson

## Abstract

**Background:** Steroid 5β-reductase (AKR1D1) plays important roles in hepatic glucocorticoid clearance and bile acid synthesis. Glucocorticoids and bile acids are potent metabolic regulators, but whether AKR1D1 controls metabolic phenotype *in vivo* is unknown.

**Methods:** *Akr1d1-/-*mice were generated on a C57BL/6 background. Liquid chromatography / mass spectrometry, metabolomic and transcriptomic approaches were used to determine effects on glucocorticoid and bile acid homeostasis. Metabolic phenotypes including body weight and composition, lipid homeostasis, glucose tolerance and insulin sensitivity were evaluated. Molecular changes were assessed by RNASeq and western blotting. *Male Akr1d1-/-*mice were challenged with a 60% high fat diet.

**Results:** *Akr1d1-/-*mice had a sex specific metabolic phenotype. At 30-weeks of age male, but not female, *Akr1d1-/-*mice were more insulin sensitive and had reduced lipid accumulation in the liver and adipose tissue, concomitant with hypertriglyceridemia and increased intramuscular triacylglycerol. This phenotype was underpinned by sexually dimorphic changes in bile acid metabolism and composition, but without overt effects on glucocorticoid action. Male *Akr1d1-/-*mice were not protected against diet induced obesity and insulin resistance.

**Conclusion:** This study shows that AKR1D1 controls bile acid homeostasis *in vivo* and that altering its activity can affect insulin sensitivity and lipid homeostasis in a sex dependent manner.

## 1 Introduction

Glucocorticoids and bile acids are potent metabolic hormones that play a critical role in the regulation of metabolism and energy balance. This is exemplified in patients with glucocorticoid excess, Cushing’s Syndrome, who develop a broad adverse metabolic phenotype^1^, and by the metabolic improvements seen in patients with steatohepatitis following treatment with bile acid sequestrants^2,3^.

Glucocorticoid availability to bind its receptor is not only dependent on circulating levels but also the tissue specific complement of pre-receptor steroid metabolizing enzymes. Best described are 11β-hydroxysteroid dehydrogenase 1 (11β-HSD1) and the 5α-reductases type 1 and 2 (5αR1 and 2). 11β-HSD1 converts the inactive glucocorticoid cortisone to its active form cortisol, and *11*β*-Hsd1-/-*mice have a beneficial metabolic phenotype with improved insulin sensitivity and protection against hepatic steatosis^4–6^. 5αRs catalyze the first step in cortisol clearance to 5α-tetrahydrocortisol and 5αR1-/-mice have increased hepatic steatosis on a western diet^7,8^, and patients treated with 5αR inhibitors have increased intrahepatic lipid accumulation^9^, skeletal muscle insulin resistance^10^ and risk of type 2 diabetes^11^.

The metabolic effects of bile acids are primarily mediated through the farnesoid X receptor (FXR) and Takeda G-protein receptor 5 (TGR5), although bile acids and intermediates of their synthesis can activate or antagonize multiple metabolic receptors, including the liver X receptor (LXR) and pregnane X receptor (PXR)^12^. The primary bile acids, cholic acid (CA), chenodeoxycholic acid (CDCA), and in mice α- and β-murocholic acid (α/β-MCA), are synthesized from cholesterol in the liver and once released into the intestine can be further metabolized by bacterial enzymes to secondary bile acids, deoxycholic acid (DCA), lithocholic acid (LCA), and ω-MCA^13^. As bile acid receptors have differing affinities for each bile acid species, metabolic consequences are not only dependent on total bile acid levels but also composition of the bile acid pool. This is highlighted by the phenotype of *Cyp8b1-/-*mice, a sterol 12α-hydroxylase required for the generation of CA. These animals have a complete absence of CA and its derivatives, and metabolically, they are more insulin sensitive and protected against diet induced obesity, a phenotype reversed by supplementation with tauro-CA^14,15^.

The enzyme Δ4-3-oxosteroid 5β-reductase, encoded by the gene *AKR1D1* (named *Akr1d1* or *Akr1d4* in mice), is the 5β-reductase for all C19-C27 steroids (which include glucocorticoids and bile acids). It plays an important role in glucocorticoid clearance where, analogous to 5αR, AKR1D1 is the first step in the clearance of cortisol to tetrahydrocortisol (5β-THF), and cortisone to tetrahydrocortisone (5β-THE). It is also an essential step in bile acid synthesis, with 5β-reduction being required for generation of both CA and CDCA^16^. Patients with loss of function mutations in AKR1D1 have altered glucocorticoid and bile acid metabolism^17^. Low urinary 5β-reduced glucocorticoids and total glucocorticoid metabolites suggest a prolonged cortisol half-life and a resetting of the HPA axis to maintain normal circulating levels. In these patients, however, urinary bile acids are almost absent, suggesting a more pronounced effect on bile acid homeostasis. These patients develop neonatal cholestasis thought to be due to an accumulation of toxic bile acid precursors and 5α-reduced (allo) bile acids, although there is evidence of spontaneous recovery. Nothing is known about their metabolic status^17^.

Despite being potentially central in the regulation of glucocorticoid and bile acid availability, the role of AKR1D1 in metabolic homeostasis is almost entirely unexplored. We have recently shown that manipulating AKR1D1 activity *in vitro* alters glucocorticoid and bile acid action with effects on insulin signaling, as well as carbohydrate and lipid metabolism^18–20^. To investigate its role *in vivo* we generated an *Akr1d1-/-*mouse.

## 2 Methods

### 2.1 Strain generation

The *Akr1d1*-/-strain was generated from targeted embryonic stem (ES) cells obtained from the KOMP repository (www.komp.org; project ID VG12494). Mice were rederived to the Medical Research Council (MRC) Harwell Mary Lyon Centre specified pathogen free facility and maintained on C57BL/6NTac. The *Akr1d1* tm1 allele (*Akr1d1*^tm1(KOMP)Vlcg^) was converted to tm1.1 by cre recombination to remove the neo cassette. *Akr1d1*^tm1/+^ mice were crossed to mice carrying a ubiquitously expressed cre (C57BL/6NTac-Tg(ACTB-cre)3Mrt/H). Offspring from this cross carrying the converted allele *Akr1d1*^tm1.1/+^, and the cre recombinase, were bred to C57BL/6NTac to remove the cre allele. *Akr1d1*^tm1.1/+^ were crossed to C57BL/6NTac to increase numbers and *Akr1d1*^tm1.1^ heterozygotes intercrossed to produce Akr1d1^tm1.1/tm1.1^ and Akr1d1^+/+^ for phenotyping. *Akr1d1*-/-showed normal Mendelian inheritance (385 mice: WT 102; Het 179: *Akr1d1*-/-104) and sex ratios (385 mice: female 189; male 196).

### 2.2 Husbandry and experimental design

*Akr1d1-/-*mice were kept and studied in accordance with UK Home Office legislation and local ethical guidelines issued by the MRC (Responsibility in the Use of Animals for Medical Research, July 1993; home office license 30/3146). All procedures were conducted in accordance with the Animals (Scientific Procedures) Act 1986 Amendment Regulations 2012 (SI 4 2012/3039) and approved by the local Animal Welfare and Ethical Review Board (AWERB). Mice were kept under controlled light (light 7am–7pm, dark 7pm–7am), temperature (21±2°C) and humidity (55±10%). They had free access to water (9–13 ppm chlorine) and standard diet (SDS Rat and Mouse No. 3 Breeding diet, RM3) until 10 weeks of age when they were transferred to a high fat (60% kcal from fat; D12492; Research Diets) or matched control diet (10% kcal from fat; D12450J; Research Diets).

Male and female cohorts were bred for longitudinal metabolic phenotyping. They were housed in single sex groups of mixed genotype across multiple litters and were not randomized into groups. Experimental groups of 15 were used, with sample size estimates based on previous experience of mouse models in which metabolic traits were measured^8,21^.

### 2.3 Metabolic assessments

Body weight was measured weekly in the morning using average weights (g) calculated by Adventure Pro balances (Ohaus, US). Fat and lean mass was assessed by Echo-MRI (Echo Medical System, Houston, US) at 10 weeks of age and body composition by high energy X-Rays using the Lunar PIXImus (GE Medical, US) at 29 weeks.

Calorimetry data was collected in a PhenoMaster system (TSE Systems, Germany) at 11 weeks of age. Data were collected at 3-4 time points each hour and measurements included photobeam activity monitoring, food intake, and indirect gas calorimetry that simultaneously measures oxygen consumption (VO_2_), carbon dioxide production (VCO_2_), and respiratory exchange ratio (RER). Fecal pellets from n=7 mice were collected over 24 hours and stored at -20°C before energy content was measured by bomb calorimetry (IKA C2000 Basic, IKA Oxford) as previously described^22^ and triacylglycerol by colorimetric assay (Cayman Chemical, Michigan, US)

To measure intraperitoneal or oral glucose tolerance (ipGTT and oGTT) n=14-15 mice were fasted overnight then either injected intraperitoneally with 20% glucose solution (2g glucose/kg body weight; Sigma, UK) or orally gavaged with 1g glucose/kg body weight. Glucose concentration was measured in the tail vein blood of restrained animals at t=0, 15, 30, 60, and 120 minutes (Alphatrak, Abbott, UK). To measure insulin tolerance (ipITT) mice were fasted for 4-5 hours then injected intraperitoneally with insulin at 0.75 IU/kg for the females and 1.25 IU/kg for males (Hypurin Bovine Insulin). Glucose concentration was measured in the tail vein blood of restrained animals at t=0, 15, 30, 45 and 90 minutes (Alphatrak, Abbott, UK).

### 2.4 Blood biochemistry and metabolomics

At termination mice were anaesthetized with isoflurane and blood collected *via* retro-orbital bleed. Samples were kept on ice then centrifuged for 10 minutes at 8,000 x g at room temperature. Corticosterone (Enzo Corticosterone ELISA Kit, Lausen, Switzerland), insulin (CrystalChem Ultra-Sensitive Mouse Insulin ELISA, Zaandam, Netherlands) and GLP-1 (CrystalChem Mouse GLP-1 ELISA, Zaadam, Netherlands) were measured by ELISA. Triacylglycerol, total cholesterol, LDL cholesterol, alanine transaminase (ALT) and aspartate transaminase (AST) were measured using Instrumentation Laboratory kits on an ILab 650 Clinical Chemistry analyzer with manufacturer recommended reagents and settings. Unbiased plasma metabolomics was performed by Metabolon (Metabolon, Inc., Research Triangle Park, NC, USA) using their global mouse metabolite panel and according to published methods^23^. Metabolon data are presented as log2(FC) with p values generated from relative signal intensity from n=10 mice.

### 2.5 LC-MS/MS quantification of bile acids and their intermediates and GC-MS quantification of sex steroids

Extraction and quantification of bile acids from plasma (25 µl) and liver tissue (30±10 mg) was performed using the protocol from Penno et al^24^ with the following modifications: Plasma samples were diluted with water (75 µl) and subjected to protein precipitation by isopropanol (900 µl, containing 100 nM internal standards). Samples were incubated at 4°C and 1400 rpm for 30 minutes and centrifuged at 4°C and 16,000 x g for 10 minutes. Liver was homogenized at 4°C and 6500 rpm for 3 x 30 seconds in water-chloroform-methanol (1 ml; 20/20/60 v/v/v containing 100 nM internal standards) on a Precellys homogenizer (Bertin Instruments; Rockville, MD, USA) and incubated at 37°C and 850 rpm for a further 15 minutes. Samples were centrifuged at room temperature and 16,000 x g for 10 minutes and 800 µl of supernatant collected. Sample extractions were repeated twice. Injection volume 2 µl for plasma and 3 µl for liver. Quantification was conducted as described in Penno et al^24^ with minor modifications: eluent gradients were set from 0-8 minutes (25%), 8-17.5 minutes (35-68%), followed by a wash out 17.5-18 minutes (68-25%), 18-20 minutes (25-100%) and 20-22 minutes (100%). Flow rate was set to 0.63 ml/min. Experimental group sizes were n=11-15 mice. Statistical analysis of relative abundance was calculated from % of total bile acids using two-tailed unpaired parametric t-tests and significance defined by a false discovery rate (Benjamini, Krieger and Yekutieli method) adjusted p-value < 1%. Principal component analysis was performed using the FactoMineR package^25^ and the factoextra package to visualise the results in R.

Free and esterified oxysterols were measured as previously described^26^ with the following modifications: Liver (100 mg) was spiked with 30 µl of 1 μM internal standard mix 25(R/S), 26-Hydroxycholesterol-d4, 7α-hydroxy-4-cholesten-3-one-d7, 7α,12α-dihydroxycholest-4-en-3-one-d7) (Toronto Research Chemicals, Ontario, Canada) and homogenized in chloroform/methanol (4 ml: CHCl3/MeOH, 2:1, v/v) containing 50 μg/mL butylated hydroxytoluene. Oxysterols were subsequently extracted by solid phase extraction using 100 mg Silica SPE columns (Waters, Hertfordshire, UK). Samples were dried under constant stream of N_2_ and reconstituted in 125 μl of methanol for analysis by LC-MS/MS. The transitions monitored were previously reported^26^ with the addition of 7α-hydroxy-4-cholesten-3-one (401.3→383.0 m/z), 7α-hydroxy-4-cholesten-3-one-d7 (408.3→390.3 m/z), 7α,12α-dihydroxycholest-4-en-3-one (417.3→381.3 m/z) and 7α,12α-dihydroxycholest-4-en-3-one-d7 (424.3→388.3 m/z). Oxysterols were quantified relative to a calibration series ranging from 0.01-2 μM and concentrations calculated relative to their deuterated internal standards. Experimental group sizes were n=10 mice.

The concentrations of serum androstenedione, testosterone and 5α-dihydrotestosterone were determined with a validated gas chromatography tandem mass spectrometry method^27^.

### 2.6 Tissue histology and biochemistry

Adipose (gonadal and subcutaneous) tissue was fixed in 4% buffered paraformaldehyde and samples subsequently paraffin-embedded and 5 μm sections prepared on a microtome (Leica). Adipose sections were viewed at 20 x magnification, and adipocyte cross-sectional area calculated using Adobe Photoshop 5.0.1 (Adobe Systems, San Jose, CA) and Image Processing Tool Kit (Reindeer Games, Gainesville, FL). Experimental group sizes were n=8 mice. The minimum number of cells required to determine cell size distribution was calculated as previously described^28^ and statistical significance assessed using a Wilcoxon signed-rank test. Tissue triacylglycerol (Cayman Chemical, Michigan) and glycogen (Biovision, Milpitas, US) were measured in snap frozen tissue using colorimetric assays.

### 2.7 RNA sequencing

Total liver RNA was extracted from n=10 mice using a RNeasy Plus mini kit (Qiagen, Hilden, Germany). Total gonadal adipose RNA was extracted using a modified Tri-reagent (Sigma-Aldrich, Dorset, UK) protocol. Tissue (50 mg) was homogenized in 1ml of ice-cold Tri-Reagent and incubated at room temperature for 5 minutes, 200 μL of chloroform was added, the sample vigorously shaken, and incubated at room temperature for 5 minutes. After centrifugation at 12,000 x g for 15 minutes the aqueous phase was combined with an equal volume of 70% ethanol, vortexed and transferred to an RNeasy Lipid Tissue spin column (Qiagen, Hilden, Germany) for washing and elution. Concentration was determined spectrophotometrically at OD260 on a Nanodrop spectrophotometer (Thermo Scientific, Hemel Hempstead, UK) and quality on an RNA Bioanalyzer chip (Agilent, Santa Clara, US).

Following extraction, RNAs were incubated with oligo (dT) beads and enriched poly-A libraries were selected using TruSeq Stranded mRNA HT Sample Prep Kit for Illumina with custom 12 bp indexes. Libraries were multiplexed (10 samples per lane), clustered using HiSeq 3000/4000 PE Cluster Kit and paired-end sequenced (75 bp) using in-house indexes to a total depth of ∼25 million read pairs, on the Illumina HiSeq4000 platform. Reads were mapped with Stampy^29^ on default settings with GRCm38/mm10 as genome reference and bam files merged using Rsamtools (v2.0). Gene level read counts for all protein-coding RNA transcripts present in refTene mm10 were quantified in a strand-specific manner using FeatureCounts from the Rsubread package (v1.34.6). Differential expression analysis was performed using EdgeR (v3.26.6)^30^ on normalized genes counts using the trimmed mean of M-values (TMM) method) for all autosomal protein-coding genes that were expressed at > 0.25 counts per million (CPM) in at least 2 samples. Statistical comparisons were performed using the glmLRT function in EdgeR and using an adjusted p-value < 0.05% (Benjamini-Hochberg method). Ingenuity Pathway Analysis (IPA, QIAGEN Redwood City, www.qiagen.com/ingeniuty) was used to predict causal networks and upstream regulators. The expression levels of key regulated genes were confirmed by qPCR (Table.S2).

### 2.8 Cell culture and treatments

Huh7 cells were a gift from Dr. Camilla Pramfalk (Karolinska Institutet). Cells were cultured in Dulbecco’s Minimum Essential Medium (DMEM) (Zen Bio Inc., Durham, NC, USA), containing 4.5□g/L glucose, and supplemented with 10% fetal bovine serum, 1% non-essential amino acids and 1% penicillin/streptomycin. Cortisol, cholic acid (CA), chenodeoxycholic acid (CDCA), and GW4064 were purchased from Sigma-Aldrich (Dorset, UK). Cells were treated for 24 h with 500 nM cortisol, 50 μM CA or 50 μM CDCA. All treatments were performed in serum-free and phenol-red free media, containing 4.5□g/L glucose (Zen Bio Inc., Durham, NC, USA).

### 2.8 Reverse transcription and quantitative PCR

Total RNA was extracted from snap frozen tissue (n=10 liver and quadricep muscle) and cells using Tri-Reagent (Sigma-Aldrich, Dorset, UK) and concentration determined spectrophotometrically at OD260 on a Nanodrop spectrophotometer (Thermo Scientific, Hemel Hempstead, UK). Reverse transcription was performed on 1 μg of RNA using the High-Capacity cDNA Reverse Transcription Kit with RNase Inhibitor (Thermo Scientific, Hemel Hempstead, UK). All quantitative PCR experiments were conducted on an ABI 7900HT (Perkin-Elmer Applied Biosystems, Warrington, UK). Reactions were performed in a 6 μl volume of 1 x Kapa Probe Fast Mastermix (Kapa Biosystems, Amsterdam, Netherlands). TaqMan assays (FAM labelled) and all reagents were supplied by Applied Biosystems (Applied Biosystems, Foster City, US). The reaction conditions were: 95°C for 3 minutes, 40 cycles of 95°C for 3 seconds and 60°C for 20 seconds. The Ct of each sample was calculated using the following equation (where E is reaction efficiency determined from a standard curve): ΔCt = E^[min Ct-sample Ct]^ using the 1/40 dilution from a standard curve generated from a pool of all cDNAs as the calibrator. Relative expression ratio was calculated using the equation: ratio = ΔCt_[target]/_ ΔCt_[ref]_ and expression normalized to the geometric mean of 18S rRNA and HPRT. Statistical analysis was performed on mean relative expression ratio values (Ratio= ΔCt[target]/ΔCt).

### 2.9 Protein extraction and immunoblotting

Total protein was extracted from snap frozen tissue and cells using RIPA buffer (Sigma-Aldrich, Dorset, UK), with protease (1/100) and phosphatase inhibitor cocktail (1/100) (ThermoFisher Scientific, Loughborough, UK) and protein concentration measured using a commercially available assay (Bio-Rad, Hemel Hempstead, UK). Primary INSRβ (#3025S), IRS1 (#2390S), total-AKT (#4685S) and mTOR (#2972S) (Cell Signaling Technology, Leiden, The Netherlands) and secondary antibodies from Dako (Agilent Technologies, Santa Clara, USA) were used at a dilution of 1/1000. Protein was run on Criterion TGX Stain-Free Precast Gels and total protein visualized using the ChemiDoc MP Imaging System (Bio-Rad, Watford, UK). Bands were visualized with Bio-Rad Clarity Western ECL (Watford, Hertfordshire, UK) on a ChemiDocXS imager (Bio-Rad, Watford, UK), quantified by densitometry and normalized to total protein from the same gel using ImageJ (National Institutes of Health). Quantifications are based on n=3-10 mice. The original Western blots used for quantification are presented in supplementary figures 3 (male) and 4 (female).

### 2.10 Statistics

Data are presented as mean ± standard error unless otherwise stated. Data analysis was performed using Graphpad Prism software (Graphpad Software Inc, La Jolla, USA). Normality was assessed using the Shapiro-Wilk test. Two-tailed, unpaired t-tests were used to compare differences in mean between genotype when assumptions of normal distribution were met with Mann-Whitney tests used on datasets with nonparametric distribution. Two-way analysis of variance (ANOVA) with Sidak corrections were used to compare means grouped by sex and genotype and repeated measure two-way ANOVA for data collected across time. Comparisons were considered statistically significant at p<0.05.

## 3 Results

### 3.1 Akr1d1 deletion does not affect glucocorticoid metabolism but decreases total bile acid levels and alters bile acid composition

*Akr1d1* deletion (Fig.1A) did not significantly alter adrenal mass (male WT 1.80±0.24 *vs*. -/- 1.76±0.13 mg; female WT 3.73±0.25 *vs*. -/- 3.16±0.38 mg) or serum corticosterone levels (the major circulating rodent glucocorticoid) (Fig.1B). Consistent with no change in glucocorticoid receptor activation in the liver, hepatic expression of the glucocorticoid regulated genes serum and glucocorticoid-regulated kinase 1 (*Sgk1*), glucocorticoid-induced leucine zipper protein-1 (*Gilz*), dual specificity phosphatase 1 (*Dusp1*) and 11βHSD1 (*Hsd11b1*) as well as the glucocorticoid metabolizing enzymes 5αR1 & 2 (*Srd5a1* & *2*), *Hsd11b1*, 3α-hydroxysteroid dehydrogenase (*3*α*hsd*) and 20α-hydroxysteroid dehydrogenase (*20*α*hsd*) were unchanged (Fig.1C). Serum levels of other steroid substrates of AKR1D1 were not significantly altered (testosterone pg/ml: females undetected, male WT 417±159 *vs*. -/- 333±170; 5α-dihydrotestosterone pg/ml: females undetected, male WT 29.7±10.9 *vs*. -/- 24.8±10.4; androstenedione pg/ml: female WT 100±41 *vs*. -/- 103±30, male WT 70±17 *vs*. -/- 59±10).

**Figure 1:**
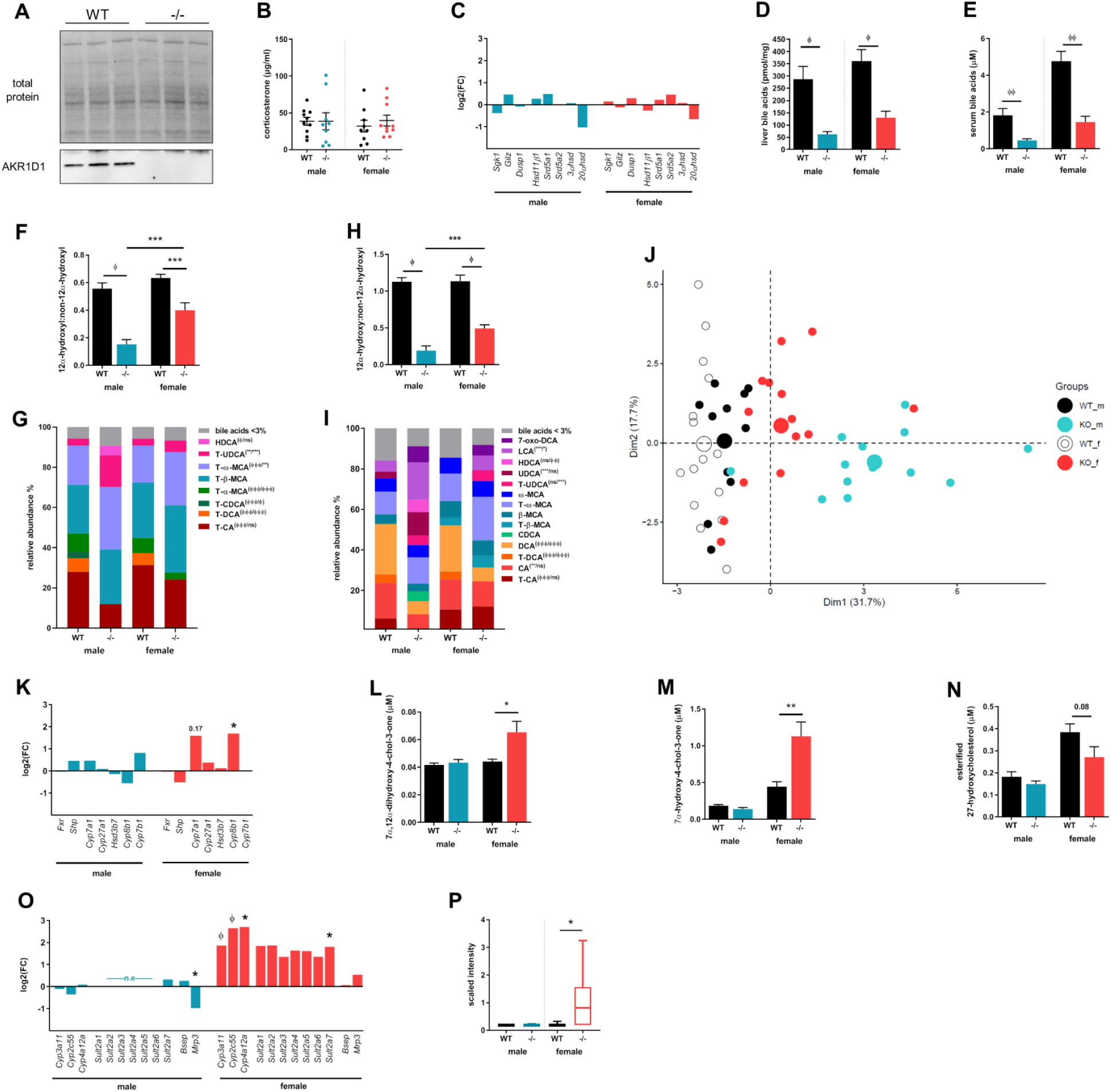
Hepatic and serum bile acids are lower in *Akr1d1-/-*mice with sexually dimorphic changes to bile acid metabolism and composition. *Akr1d1* deletion (A: western blot, liver), does not alter serum corticosterone levels (B) (n = 10 mice), or hepatic mRNA expression of glucocorticoid responsive genes, *Sgk1, Gilz, Dusp1* and *11*β*hsd1* (C) (n = 10 mice). Male and female *Akr1d1-/-*mice have decreased total hepatic (D) and serum (E) bile acids and altered bile acid composition with reduced 12α-hydroxylated/non-12α-hydroxylated bile ratio in the liver (F & G) and serum (H & I) (n = 12-15 mice). Principle Component Analysis shows greater divergence from wildtype in male *Akr1d1-/-*mice (J). *Akr1d1* deletion has a sexually dimorphic effect on mRNA expression of hepatic bile acid metabolizing genes and levels of bile acid intermediates. *Cyp8b1* expression is increased in *Akr1d1-/-*females but not males (K) (n = 10 mice) as are the AKR1D1 substrates 7α,12α-dihydroxy-4-chol-3-one (L) and 7α-hydroxy-4-chol-3-one (M) (n = 10 mice). The oxysterol 27-hydroxycholesterol (27-OHC) is decreased in *Akr1d1-/-*females (N) (n = 10 mice). Female *Akr1d1-/-*mice also have increased expression of the bile acid detoxifying genes *Cyp3a11, Cyp2c55, Cyp4a12a* and *Sult2a7* (O) (n = 10 mice) and serum levels of LCA sulfate (P) (n = 10 mice). Data are presented as mean±se, log2(FC), ratio, or mean relative abundance. *p<0.05, * *p<0.01, * * *p<0.005, ^Ø^p<0.001, ^ØØ^p< 0.0005, ^ØØØ^p<0.0001 compared to wildtype. mRNA expression was measured by RNASeq. p-values for bile acid composition compare WT and *Akr1d1-/-*within sex (male/female). (WT = wildtype C57BL/6; -/-= *Akr1d1-/-*)

Contrasting the lack of effect on glucocorticoid action and metabolism, the impact on bile acid homeostasis was marked. Total liver (Fig.1D) and serum (Fig.1E) bile acid levels were reduced, and composition altered, with a decreased 12α-hydroxylated (CA and CA-derived)/non-12α-hydroxylated (CDCA and CDCA-derived) ratio in both the liver (Fig.1F & G) and serum (Fig.1H & I). The relative reduction in 12α-hydroxylated bile acids was greater (Fig.1F & 1H) and bile acid profiles more markedly different in male *Akr1d1-/-*mice (Fig.1J). Absolute levels of liver and serum bile acids and bile acid intermediates are presented in supplementary table 1. Bile acids inhibit their own synthesis *via* FXR activation of small heterodimer partner (SHP) repressing expression of bile acid synthesizing enzymes. Despite lower hepatic bile acids there was no reduction in *Shp* expression, and in *Akr1d1-/-*males the expression of bile acid synthesizing enzymes was unchanged (Fig.1K). In females the expression of *Cyp7a1* and *Cyp8b1* were increased, with only the latter reaching significance (Fig.1K). Further suggesting an increase in *Cyp7a1* activity, intermediates of the classic pathway 7α-12α-dihydroxy-4-chol-3-one (Fig.1L) and 7α-hydroxy-4-chol-3-one (Fig.1M) were increased and there was a trend toward decreased 27-hydroxycholesterol levels (Fig.1N), the first metabolite in the alternative pathway. The bile acid synthesis pathway is presented in figure 2. In addition to altering synthesis, the expression of genes in the bile acid detoxification pathway were increased in *Akr1d1-/-*females, including the phase I (oxidation) genes *Cyp3a11, Cyp2c55* and *Cyp4a12a* as well as the phase II (conjugation) gene *Sult2a7* (Fig.1O). Expression of key regulated genes were confirmed by qPCR (Table.S2). Sulfated bile acid species were not measured by LC-MS, but consistent with increased bile acid detoxification and clearance in *Akr1d1-/-*females, serum T-lithocholate-3-sulfate was increased in a metabolomic screen (Fig.1P). Consistent with this gene expression pattern in females Ingenuity Pathway Analysis (upstream regulators) predicted activation of key transcriptional regulators of cholesterol metabolism, constitutive androstane receptor (CAR: *Nr1i3*) and PXR (*Nr1i2*) (Table.1A).

**Table 1.**
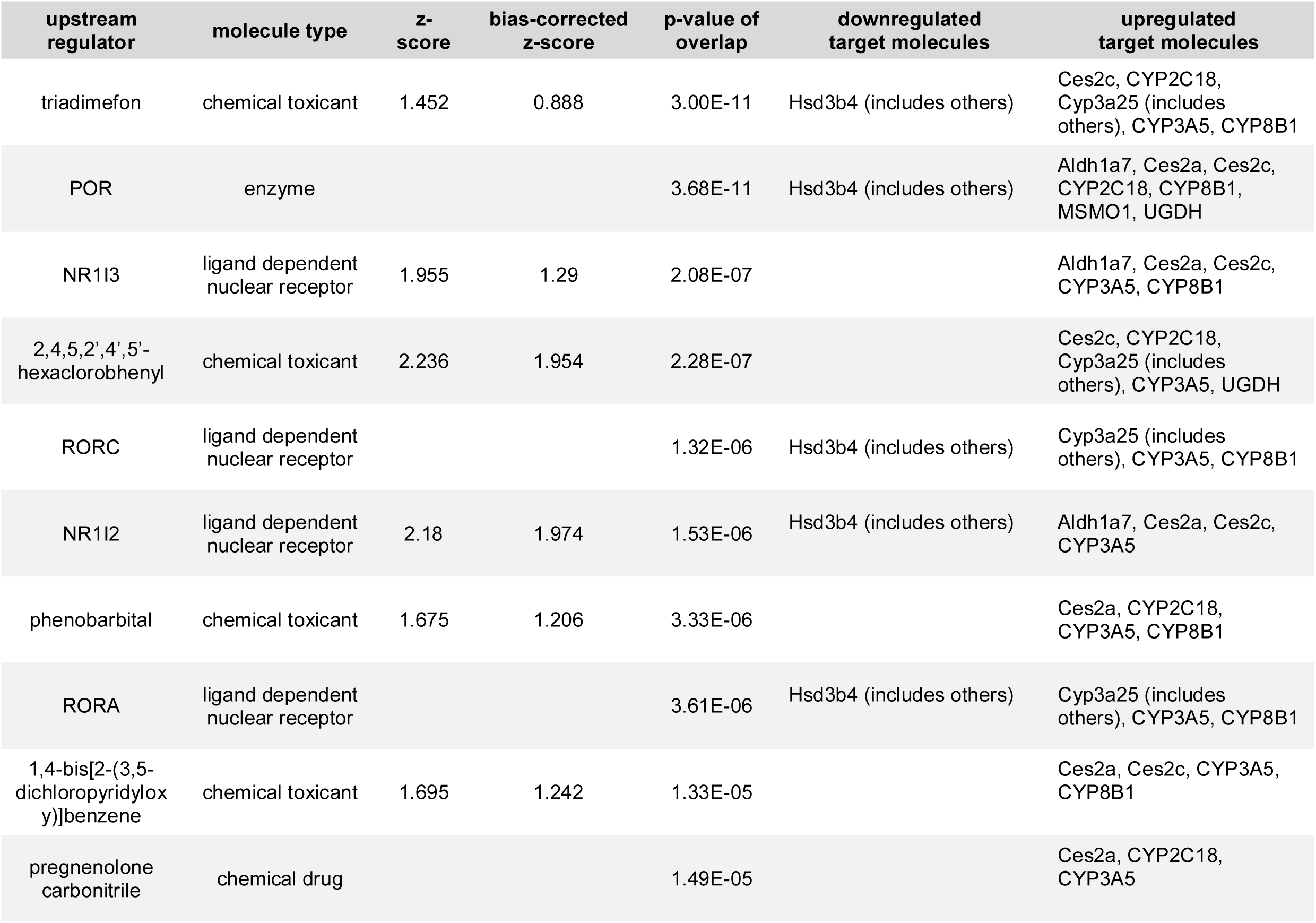
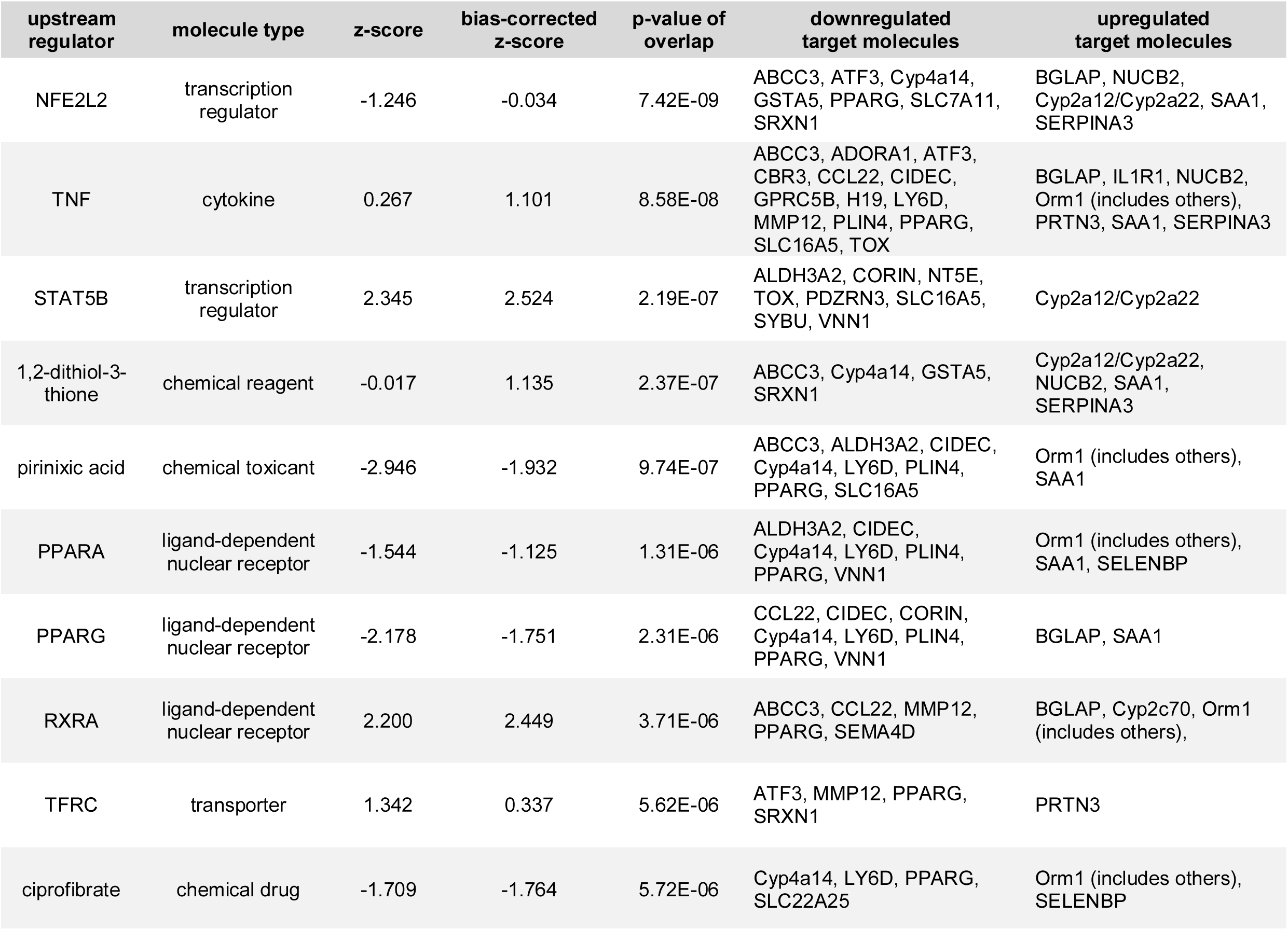
Top 10 upstream regulators in *Akr1d1-/-*liver predicted by Ingenuity Pathway Analysis. Ingenuity Pathway Analysis (IPA) of liver RNASeq identified core metabolic transcription factors as upstream regulators of the hepatic response to *Akr1d1* deletion. In females IPA predicted activation of PXR and CAR signaling (A). In males IPA predicted activation of STAT5B and RXRα signaling and inhibition of PPARα and PPARγ signaling (B). Activation z-score infers activation status of predicted regulators. Overlap p-value measures overlap between the dataset genes and genes known to be regulated by the transcriptional regulator. Analysis was performed on RNASeq data from n = 10 mice.

**Figure 2:**
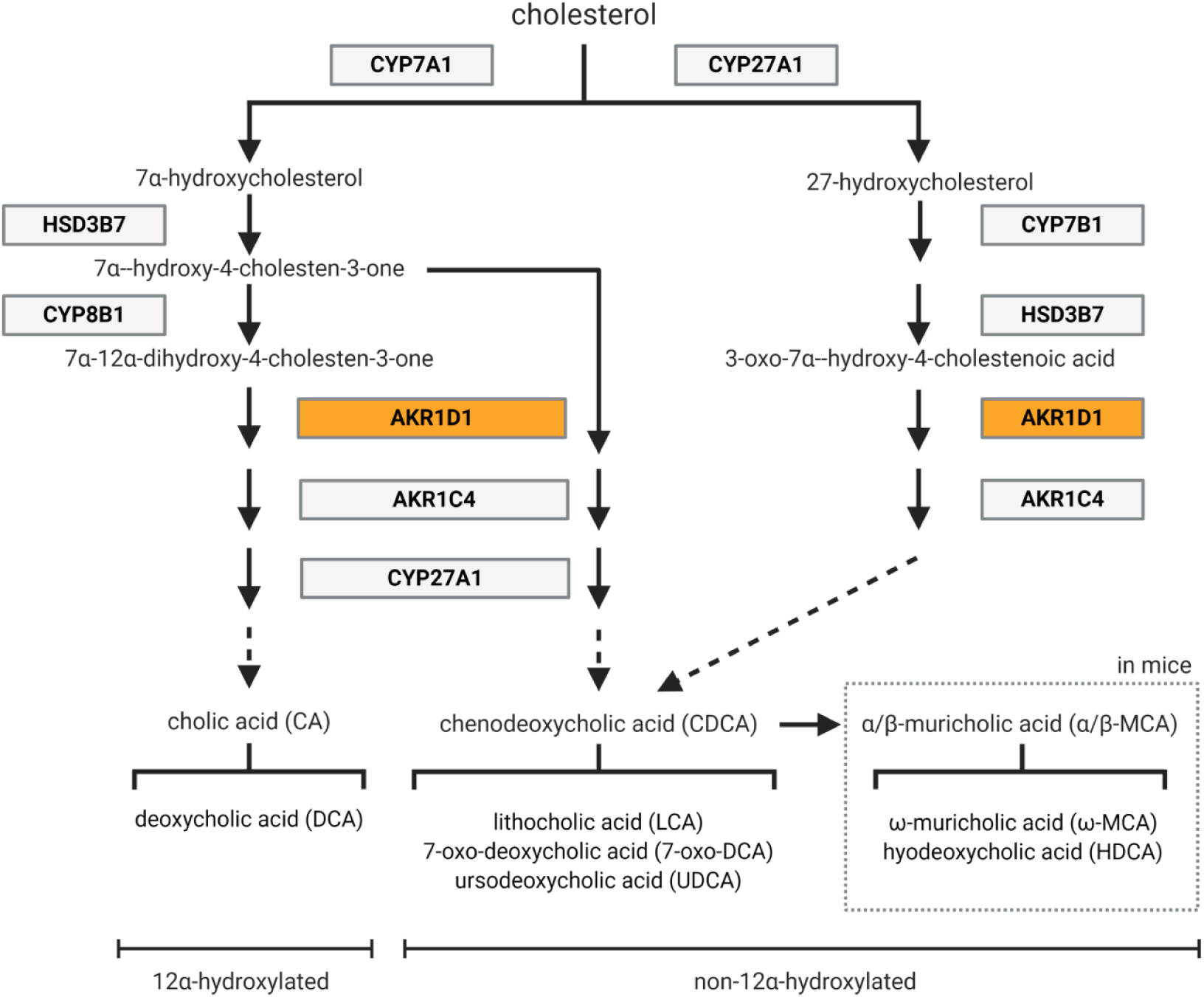
Schematic overview of the steps involved in the classic and alternative bile acid synthesis pathways in humans and mice. Figure produced in BioRender.

In contrast to patients with AKR1D1 deficiency *Akr1d1-/-*mice did not show overt signs of cholestasis, H&E (Fig.S1A), hepatic inflammation (Fig.S1B) or liver damage (Fig.S1C & D).

### 3.2 Mature (30 week) Akr1d1-/-males, but not females, have reduced fat mass and increased insulin sensitivity

Metabolic assessments were undertaken in males and females when young (10 weeks) and at maturity (30 weeks). At 10 weeks the body weight and composition of *Akr1d1-/-*mice were comparable to that of WT littermates (Fig.3A). Energy expenditure (Fig.3B) and activity rates were unchanged (Fig.S2A) but there were sex-specific effects on energy intake as well as macronutrient preference for oxidation. Male, but not female, *Akr1d1-/-*mice had increased food intake in the dark phase (Fig.3C), and a higher respiratory exchange ratio (RER), suggesting increased dependency on carbohydrates over lipids as an energy source (Fig.3D). Fecal energy (Fig.S2B) and lipid content (Fig.S2C) were normal, suggesting no increase in energy loss through malabsorption. At 10 weeks glucose control was also normal, with no change in insulin tolerance (Fig.S2D), ip or oral glucose tolerance (Fig.S2E & Fig.S2F), serum GLP-1 15 minutes post-oral glucose bolus (Fig.S2G) or in serum insulin 60 minutes post ip glucose (Fig.S2H).

**Figure 3:**
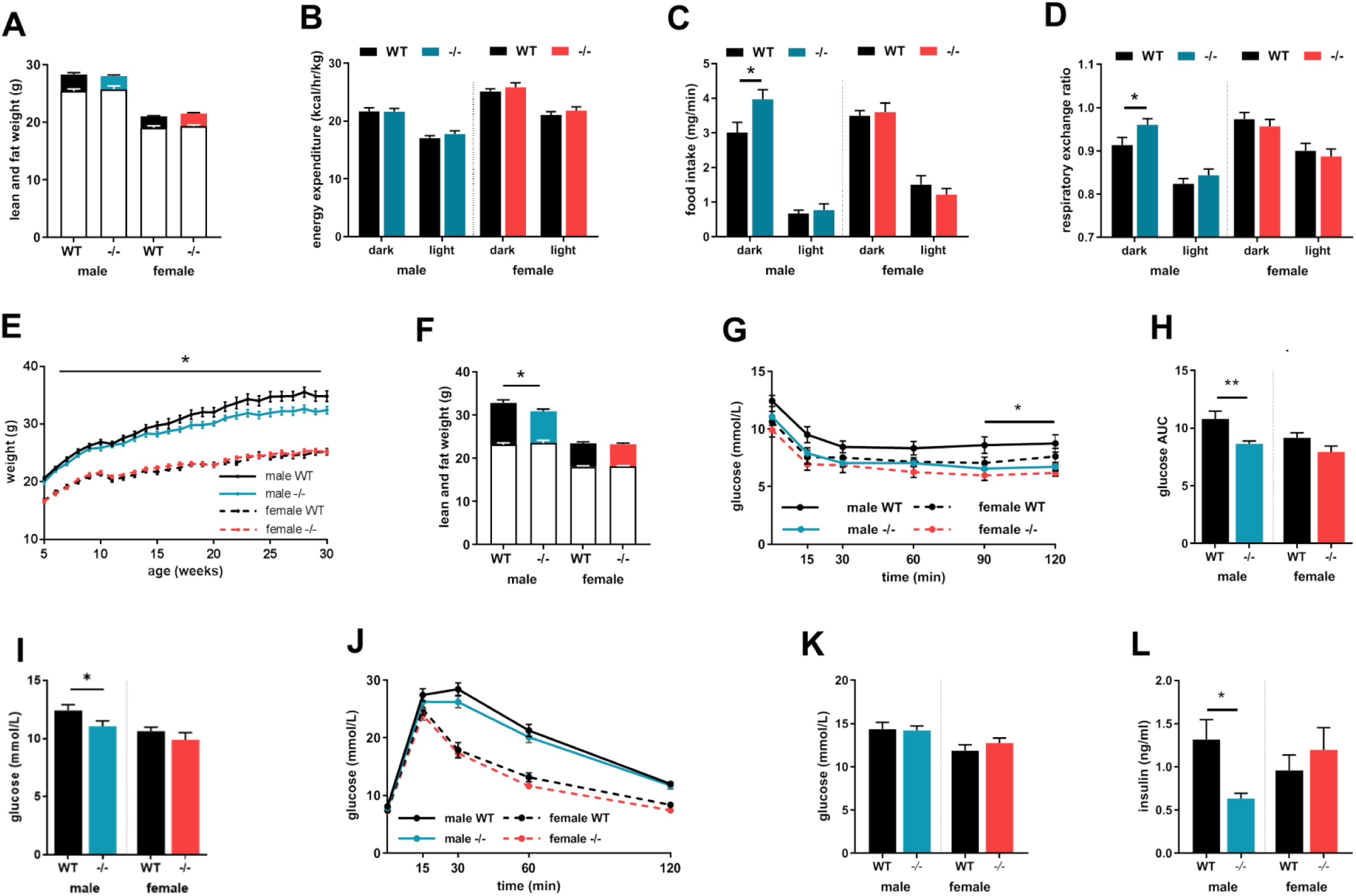
At maturity male *Akr1d1-/-*mice have reduced fat mass and increased insulin sensitivity. Young (10-week) *Akr1d1-/-*mice have normal body weight and composition (lean mass empty bar; fat mass filled bar) (A) and locomotor activity (B) is unchanged. Male, but not female, *Akr1d1-/-*mice have a preference for carbohydrate over lipid as an energy source, measured as an increase in respiratory exchange ratio (C) and increased dark phase food intake (D). Male (blue lines), but not female (red lines), *Akr1d1-/-*mice gain less weight than their WT (black line) littermates (E) and at 30 weeks have lower fat mass without change in lean mass (F) (lean mass empty bar; fat mass filled bar). *Akr1d1-/-*males have exaggerated glucose clearance after ip insulin injection (G) and as area under the curve (H). Although ip glucose tolerance (I) and fed serum glucose (J) are comparable to WT animals fed insulin concentration is decreased (K). Data are presented as mean±se of n=14-15 mice. *p<0.05, * *p<0.01 compared to wildtype of the same sex. (WT = wildtype C57BL/6; -/- = *Akr1d1-/-*)

In male mice weights diverged with age, with *Akr1d1-/-*males gaining weight at a slower rate than WT littermates (Fig.3E) and at 30 weeks DEXA body composition analysis showed decreased fat mass without change in lean mass (Fig.3F). In contrast to the 10-week cohort, mature male *Akr1d1-/-*mice had enhanced insulin sensitivity as measured across an insulin tolerance test (Fig.3G) and as area under the curve (AUC) (Fig.3H). Consistent with this finding, fasting glucose was reduced in *Akr1d1-/-*males in response to a 4 h fast (Fig.3I), however not after an 18h overnight fast (male WT 8.11±0.27 *vs*. -/-7.87±0.26 mmol/L; female WT 7.42±0.24 *vs*. -/- 7.31±0.23 mmol/L). Despite increased insulin sensitivity, ip glucose tolerance was unchanged (Fig.3J) as was fed blood glucose (Fig.3K). Circulating insulin levels were reduced in fed *Akr1d1-/-*males (Fig.3L), suggesting compensatory reduction in insulin secretion. In contrast to male mice, *Akr1d1-/-*females gained weight at the same rate as WT littermates (Fig.3E), had normal body composition (Fig.3F), insulin sensitivity (Fig.3G & 3H) and ip glucose tolerance (Fig.3I), as well as fed serum glucose and insulin (Fig.3J & Fig.3K).

At a molecular level and consistent with insulin sensitivity, quadricep muscle insulin receptor subunit β (INSRβ), total-AKT, and mTOR protein expression were increased in male *Akr1d1-/-*mice (Fig.4A & 4B). The mRNA expression of insulin receptor substrate 1 (*Irs1)* was increased and *Insr*β decreased, Pi3-kinase regulatory subunit, *Akt1* and *Mtor* expression were unchanged (Table.S2). Muscle glycogen, a product of anabolic metabolism was increased (Fig.4C). Hepatic total-AKT protein expression was increased (Fig.4D & 4E), but INSRβ, mTOR protein levels, the mRNA expression of key insulin signaling components (Table.S2) and glycogen levels were unchanged (Fig.4F). The protein expression of INSRβ, IRS1, total-AKT and mTOR were also unchanged in the gonadal fat (Fig.S3). In female *Akr1d1-/-*mice protein expression of total-AKT was increased in the quadricep muscle and liver, however, consistent with no change in global insulin sensitivity, INSRβ levels were decreased (Fig.S4) and skeletal muscle and hepatic glycogen levels were normal (Fig.S4C & F).

**Figure 4:**
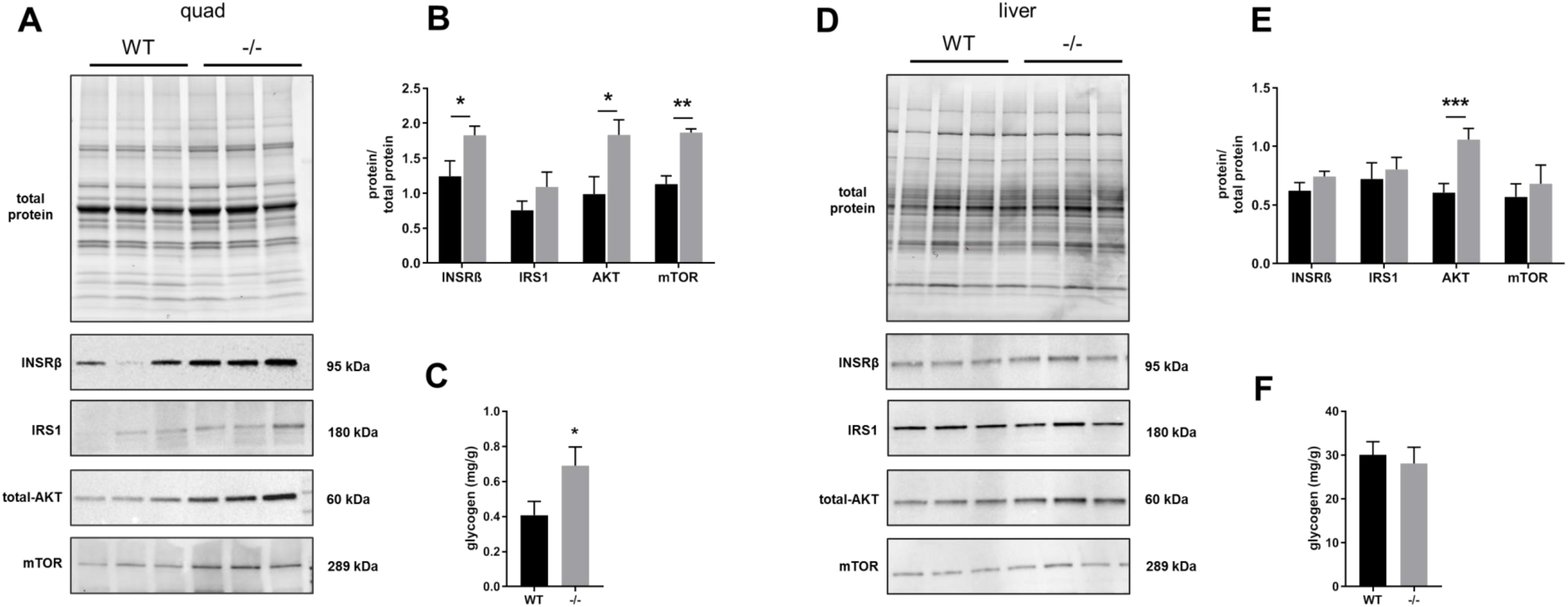
Male *Akr1d1-/-*mice have higher expression of insulin signaling components. Male *Akr1d1-/-*mice (grey bars) have increased quadricep muscle INSRβ, total-AKT, and mTOR protein expression (A & B) (quantification on n = 3-6 mice). Quadricep glycogen levels are increased (C) (n = 10 mice). In the liver, protein expression of total-AKT is increased (D & E) (quantification on n = 6 mice) and hepatic glycogen levels are normal (F) (n = 14-15 mice). Representative western blots are shown, blots used for quantification are presented in supplementary figures 3 and 4. Data are presented as mean±se. *p<0.05 compared to wildtype. (WT = wildtype C57BL/6; -/- = *Akr1d1-/-*)

Demonstrating that changes in bile acid composition have the potential to alter insulin sensitivity CDCA but not CA reduced AKT1 mRNA expression in Huh7 hepatoma cells (Fig.5A). AKR1D1 knockdown (Fig.5B) reduced media CA and CDCA levels (Fig.5C), and when calculated as a change in bile acid levels CDCA was reduced to a greater degree than CA (Fig.5D). Mirroring that seen in male *Akr1d1*-/-mice AKR1D1 knockdown increased AKT1 gene expression (Fig.5E) and protein levels (Fig.5F). Furthermore, CDCA, but not CA, was able to partially recover the increase in AKT1 expression seen in knockdown cells (Fig.5G).

**Figure 5:**
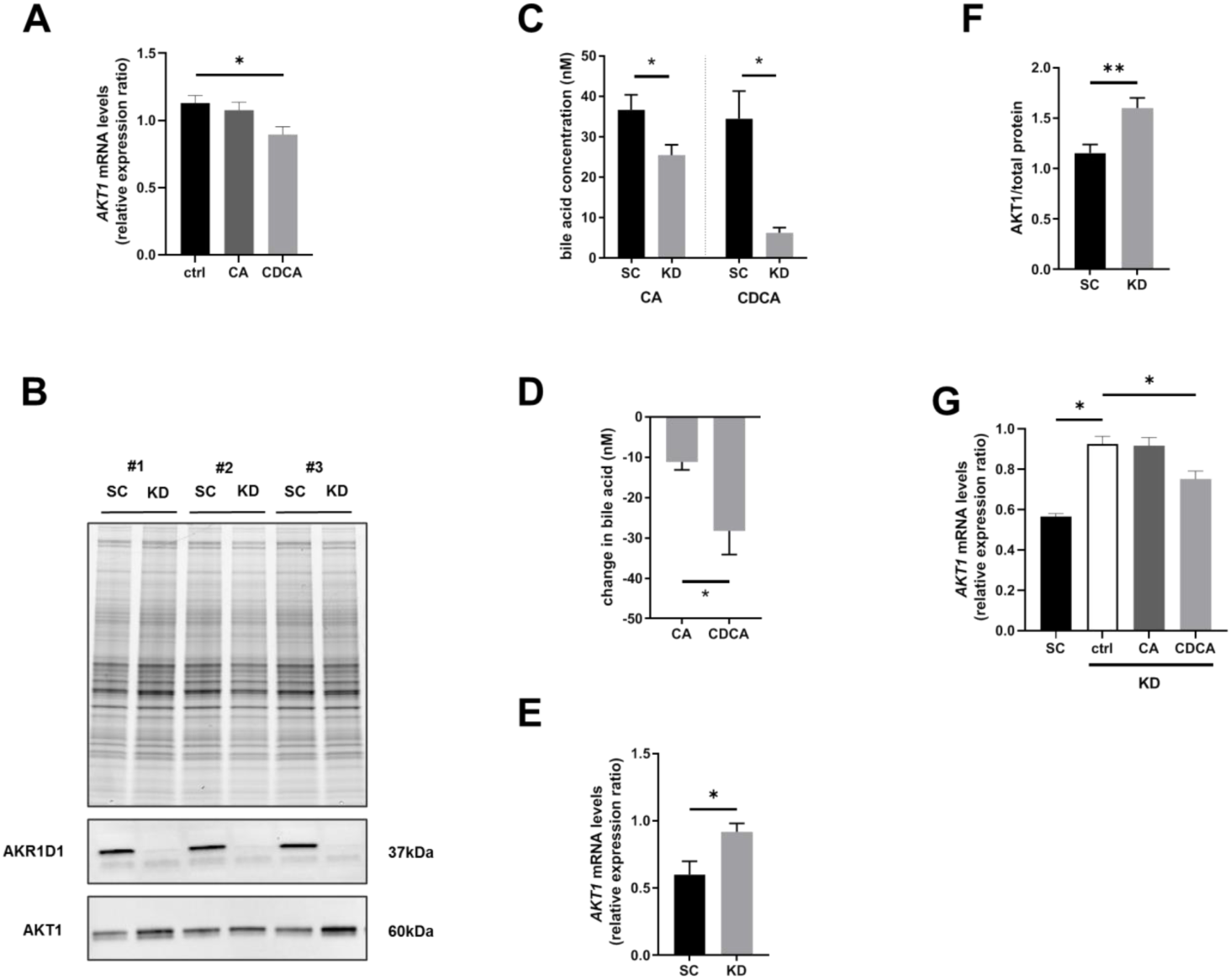
AKR1D1 knockdown increases AKT1 expression and is partially recovered by CDCA but not CA. Chenodeoxycholic acid (CDCA) but not cholic acid (CA) deceases *AKT1* gene expression (A) in Huh7 hepatoma cells. AKR1D1 deletion (B: Western blot) reduced media CA and CDCA levels (C) with CDCA reduced to a greater degree than CA (D). AKT1 gene expression (E) and protein levels (F) were increased after AKR1D1 knockdown and was partially recovered by treatment with CDCA but not CA (G). Data are presented as mean±se and n = 3-5. *p<0.05 compared to wildtype. (SC = scrambled control, KD = knockdown).

### 3.3 Akr1d1-/-males have reduced hepatic and adipose lipid stores and hypertriglyceridemia

Male *Akr1d1-/-*mice had reduced white adipose depot weights (Fig.6A) and adipocytes were smaller in the gonadal and subcutaneous depots (Fig.6B), suggesting a reduction in lipid accumulation rather than adipocyte number. Frequency distribution of gonadal (Fig.6C) and subcutaneous (Fig.6D) adipocytes are presented. Furthermore, hepatic triacyclglycerol accumulation was reduced in *Akr1d1-/-*males (Fig.6E). Consistent with reduced hepatic lipid storage, *Akr1d1-/-*males had increased serum triacyclglycerols (Fig.6F), monoacylglycerols and diacylglycerols (Fig.6G), and non-esterified fatty acids (Fig.6H), but without change in total or high-density-lipoprotein (HDL) cholesterol (Fig.6I). Relative intensity values for monoacylglycerols, diacylglycerols and fatty acids are presented in supplementary table 3. Hypertriglyceridemia is commonly associated with intramyocellular triacylglycerol accumulation and skeletal muscle triacylglycerol levels were increased in the *Akr1d1-/-*males (Fig.6J).

**Figure 6:**
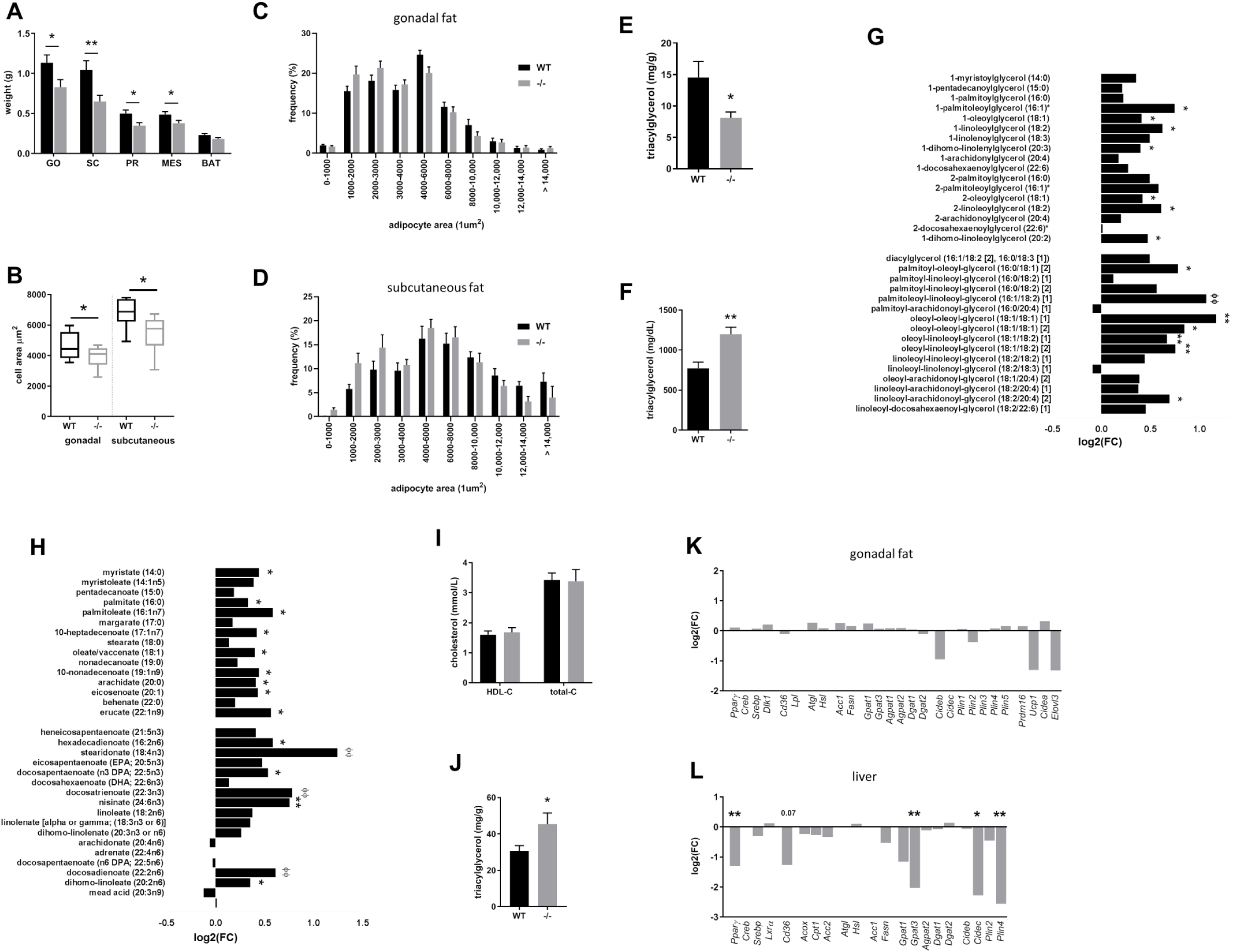
Male *Akr1d1-/-*mice have reduced adipose and hepatic lipid accumulation and hypertriglyceridemia. Male *Akr1d1-/-*mice (grey bars) have smaller white adipose tissue weights compared to wildtype littermates (black bars) (A) (n = 14-15 mice) with smaller adipocytes in the gonadal and subcutaneous depots (B: mean area; C & D: frequency distribution) (n = 8 mice) and reduced hepatic triacylglycerol (E) (n = 14-15 mice). Serum triacylglycerol (F), monoacylglycerols and diacylglycerols (G), and non-esterified fatty acids (H) are increased but total and high-density lipoprotein cholesterol (I) are normal (n = 10 mice). Intra-muscular triacylglycerol is increased in *Akr1d1-/-*quadricep muscle (J) (n = 9 mice). The mRNA expression of lipid metabolism genes in the gonadal fat are unchanged (K) but in the liver the expression of fatty acid esterification (*Gpat3*) and lipid droplet (*Cidec* & *Plin4*) genes, as well as the transcription factor *Ppar*γ are reduced (L) (n = 10 mice). *p<0.05, * *p<0.01, * * *p<0.005, ^Ø^p<0.001, ^ØØ^p<0.0005 compared to wildtype. (WT = wildtype C57BL/6; -/- = *Akr1d1-/-*). mRNA expression was measured by RNASeq.

Despite hypertriglyceridemia and reduced adipose mass, there was no change in the expression of key lipid metabolism genes in the gonadal fat from *Akr1d1-/-*males (Fig.6K) and Ingenuity Pathway Analysis (causal network) did not predict altered lipid accumulation or adipose browning. In the liver, genes involved in fatty acid uptake (*Cd36*, p=0.07), esterification (*Gpat3)* and lipid storage (*Cidec* & *Plin4)* (Fig.6L) were decreased, and Ingenuity Pathway Analysis (causal network) predicted reduced lipid accumulation. Consistent with reduced lipid accumulation, Ingenuity Pathway Analysis (upstream regulators) predicted PPARγ inhibition and STAT5B activation, as well as retinoid X receptor α (RXRα) activation (Table.1B). The expression of key lipid metabolism genes in the quadricep muscle were unchanged (Table.S2).

Although in *Akr1d1-/-*females total fat mass was unchanged (Fig.3B) adipose depots were smaller (Fig.S5A), though to a lesser degree than in males. Serum total and HDL cholesterol (Fig.S5B), total serum triacylglycerol (Fig.S5C), diacylglycerol and monoacylglycerol (Fig.S5D) were all normal, but levels of some non-esterified fatty acids were reduced (Fig.S5E). Hepatic triacyclglycerol (Fig.S5F) content was unchanged. Relative intensity values for monoacylglycerols, diacylglycerols and fatty acids are presented in supplementary table 3. The expression of key lipid metabolism genes were not altered in the gonadal fat (Fig.S5G) or liver (Fig.SH).

### 3.4 Male Akr1d1-/-mice are not protected against diet-induced obesity or insulin resistance

To investigate the interaction between genotype and diet, 10-week-old male mice were challenged with a 60% high fat diet (HFD) for 20 weeks. Figures include WT control data to allow comparison. On HFD, *Akr1d1-/-*males gained weight at the same rate as their WT littermates (Fig.7A), and body composition (Fig.7B), adipose weights (Fig.7C), hepatic triacylglycerol (Fig.7D), and total and HDL cholesterol (Fig.7E) were unchanged. Male *Akr1d1-/-*mice were partially protected against diet induced hypertriglyceridemia (Fig.7F) but not glucose intolerance (Fig.7G) or insulin resistance (Fig.7H).

**Figure 7:**
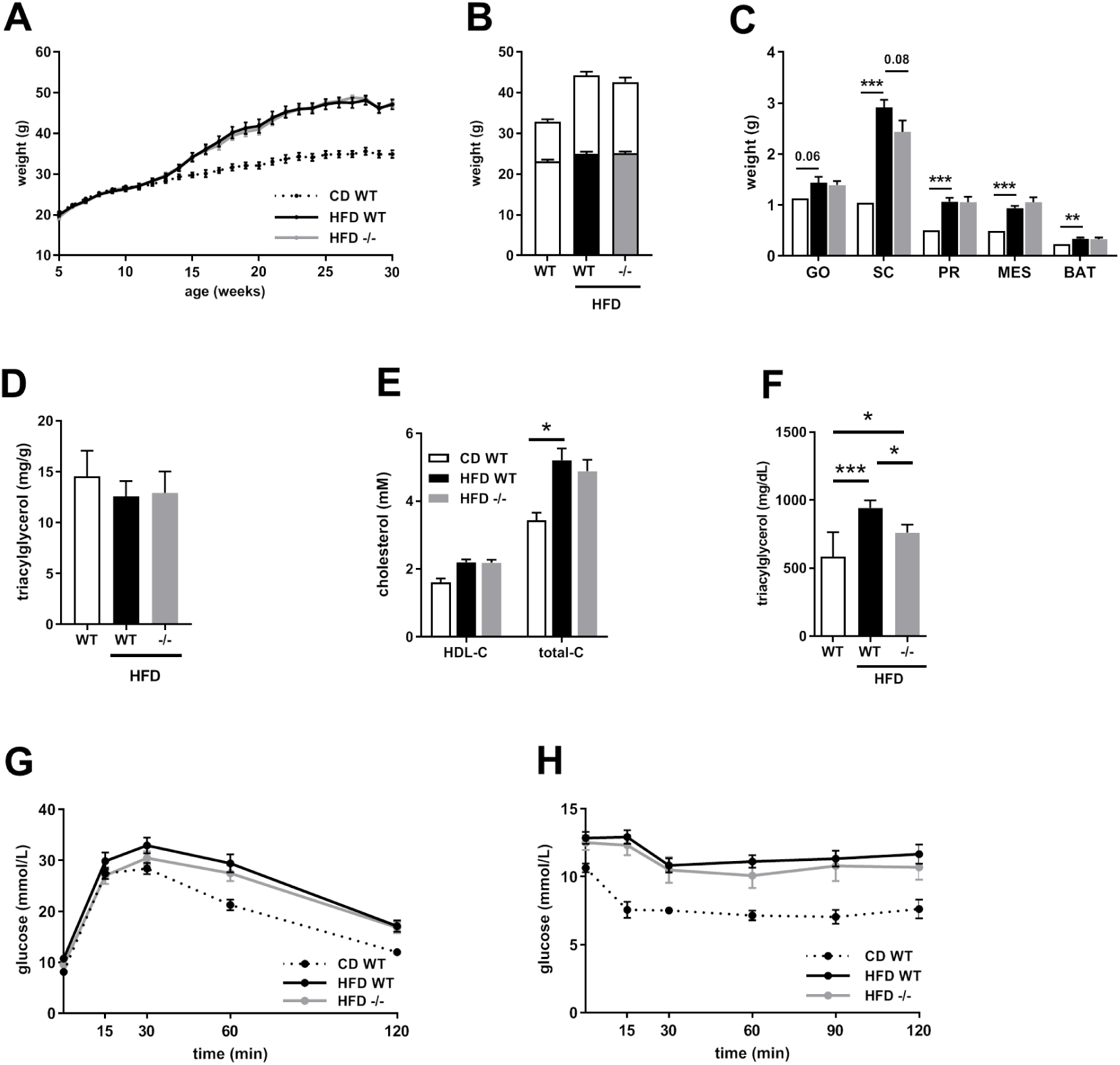
*Akr1d1* deletion does not protect male C57BL/6 mice against diet-induced obesity or its associated metabolic consequences. On a 60% high fat diet, male *Akr1d1-/-*mice (grey line) gain weight at the same rate as their WT littermates (black line) (A) and after 20 weeks body composition is not different between genotypes (lean mass lower bar; fat mass upper bar) (B) (n = 14-15 mice). Adipose weight (C), hepatic triacylglycerol (D), HDL and total cholesterol (E) were unchanged between HFD fed WT and *Akr1d1-/-*males. Serum triacylglycerol was reduced in HFD fed Akr1d1-/-males but not to levels seen in the control diet (F) (n = 10 mice). *Akr1d1-/-*males were not protected against diet induced reduction in ip glucose tolerance (G) or ip insulin tolerance (H) (n = 14-15 mice). Data are presented as mean±se. *p<0.05, * *p<0.01, * * *p<0.005 compared to wildtype. (WT = wildtype C57BL/6; -/- = *Akr1d1-/-*)

Total hepatic bile acids were reduced in WT mice on HFD but trended toward an increase in the serum (Fig.8A & B). *Akr1d1* deletion reduced total bile acids in both liver (Fig.8A) and serum (Fig.8B). Bile acid composition was altered (liver: Fig.8C; serum: Fig.8D) and the 12α-hydroxylated/non-12α-hydroxylated bile acid ratio reduced (liver: Fig.8E; serum: Fig.8F). Absolute levels of liver and serum bile acids are presented in supplementary table 4.

**Figure 8:**
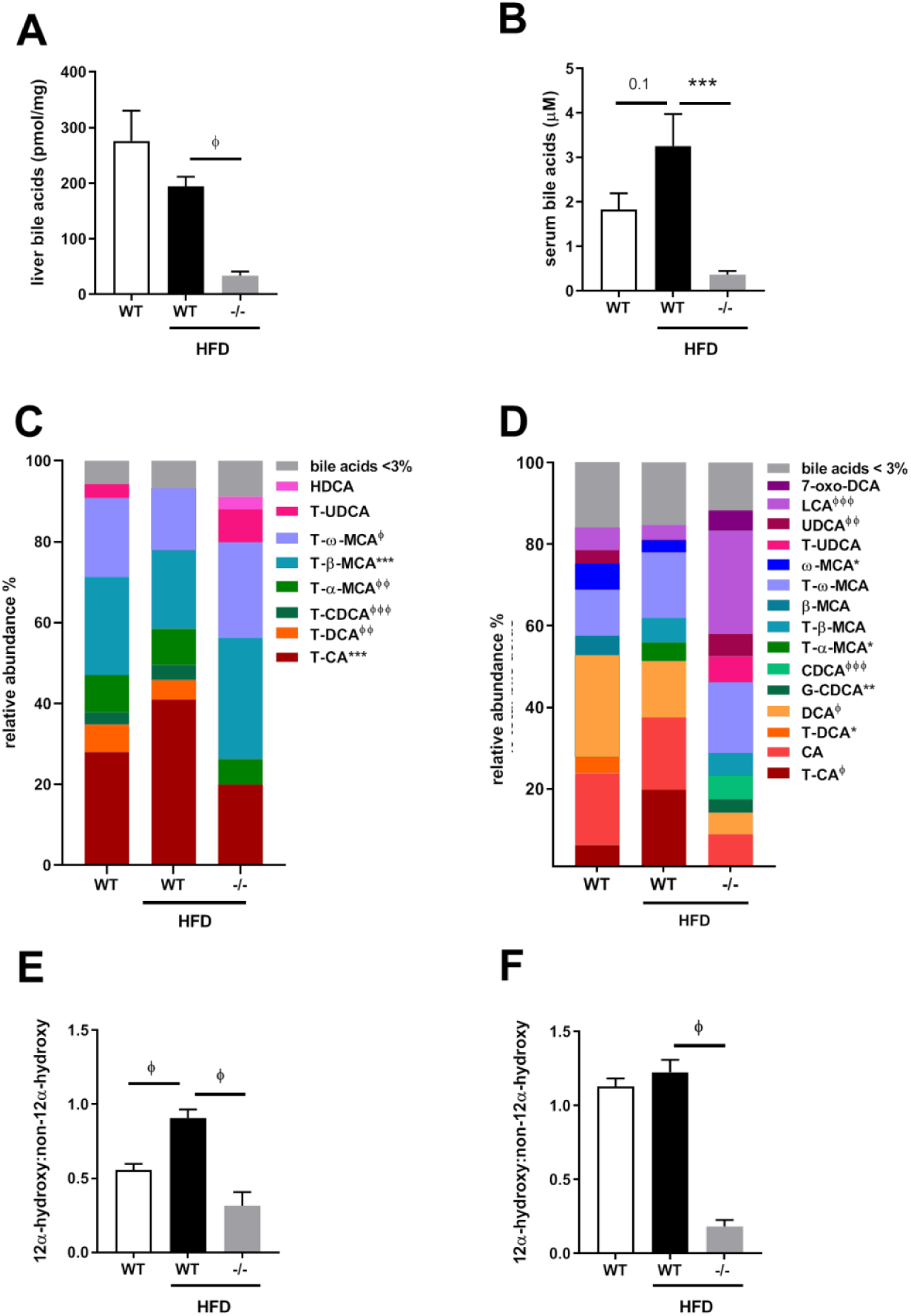
Bile acid profile in male *Akr1d1-/-*mice on HFD. Male *Akr1d1-/-*mice on HFD have reduced total liver (A) and serum (B) bile acids and altered bile acid composition (liver: C; serum D) compared to high fat fed WT littermates. Serum 12α-hydroxylated/non-12α-hydroxylated bile ratio in the liver (E) and serum (F) is reduced compared to control and HFD fed WT littermates. Data are presented as mean±se in n=10-15 mice. *p<0.05, * *p<0.01, * * *p<0.005, ^Ø^p<0.001, ^ØØ^p< 0.0005, ^ØØØ^p<0.0001. WT control diet data is presented for comparison, p value for bile acid composition compares WT and *Akr1d1-/-*on HFD. (WT = wildtype C57BL/6; -/- = *Akr1d1-/-*)

## 4 Discussion

Here we present the first *in vivo* evidence that AKR1D1 regulates metabolism, demonstrating a sex-specific metabolic phenotype in *Akr1d1-/-*mice where males, but not females, have increased insulin sensitivity and altered lipid homeostasis. Underpinning these effects are sexually dimorphic changes in bile acid metabolism and composition of the bile acid pool.

Sitting at the interface of steroid hormone and bile acid metabolism, AKR1D1 has the potential to affect metabolic homeostasis by altering steroid hormone and/or bile acid availability. Despite its central position, only a small number of studies have investigated AKR1D1 in the context of metabolic disease; hepatic gene expression is decreased in patients with type 2 diabetes^31^ and non-alcoholic fatty liver disease (NAFLD)^19^ although in one study systemic 5β-reductase activity was increased in patients with hepatic steatosis^32^. Whether reduced AKR1D1 activity contributes to the pathogenesis of metabolic disease is almost entirely unexplored, although we have recently shown that manipulating AKR1D1 alters glucocorticoid and bile acid regulation of metabolism and inflammation *in vitro*^18–20^.

AKR1D1 is the only known 5β-reductase for C19-C27 steroids, and patients with missense mutations in AKR1D1 produce only trace amounts of 5β-reduced bile acids^17^. In contrast, *Akr1d1-/-*mice still produce 5β-reduced bile acids, albeit at a lower level. This suggests the possibility of a second, yet unknown, 5β-reductase in mice and that the phenotype of the *Akr1d1-/-*mouse represents partial 5β-reductase deficiency. Patients with AKR1D1 deficiency develop severe hepatic cholestasis^33^, however we saw no evidence of cholestasis or liver damage in *Akr1d1-/-*mice. This is likely to be due to species-specific differences. During cholestasis, damage is caused by the accumulation of hydrophobic bile acids, in particular glyco-CDCA^34^, whereas mice are protected from against intrahepatic cholestasis because CDCA is converted to hydrophilic MCA species^35^.

The mechanisms underpinning the metabolic phenotype of the *Akr1d1-/-*mouse are complex but do not appear to reflect glucocorticoid excess. Mice do generate 5β-reduced 11-dihydrocorticosterone and corticosterone metabolites^36,37^ and AKR1D1 controls glucocorticoid availability and action in human hepatoma cell lines^18^, however, circulating corticosterone levels are normal in *Akr1d1-/-*mice and the hepatic expression of glucocorticoid regulated genes were unchanged. Furthermore, reduced hepatic triacylglycerol and enhanced insulin sensitivity are in direct contrast to the hepatic steatosis and insulin resistance observed in comparative models of tissue specific glucocorticoid excess (5αR1 deletion and hepatic 11βHSD1 overexpression)^7,8,21^. This lack of apparent effect on glucocorticoid action may reflect a resetting of the HPA axis to maintain circulating corticosterone levels or highlight species specific differences in steroid metabolism. In contrast to humans, where almost all excreted glucocorticoid metabolites are A-ring reduced^9^, in mice around 50% retain their 3-oxo-4-ene structure^36,38^. However, this study was designed to look at the long-term consequences of AKR1D1 deletion on metabolic health rather than as a detailed investigation into HPA axis regulation. Dedicated studies investigating the impact of AKR1D1 deletion on steroid metabolism, the stress response and glucocorticoid awakening response are needed.

The ability to activate (or antagonize) bile acid receptors differs across bile acid species^39^, making the composition of the bile acid pool of critical importance. An increase in the serum ratio of 12α-hydroxylated to non-12α-hydroxylated bile acids is associated with insulin resistance^40^. Moreover, rodent studies suggest this relationship is independent of total bile acid levels. *Cyp8b1-/-* and *Cyp7a1-/-*mice have respectively high and low total bile acids, but both have a reduced 12α-hydroxylated/non-12α-hydroxylated ratio and improved glucose control recoverable by supplementation with CA^14,41^. *Akr1d1-/-*mice have low liver and serum bile acids with a reduced 12α-hydroxylated/non-12α-hydroxylated ratio, mirroring that seen in male (females not studied) *Cyp7a1-/-*mice^41^; similarly, male *Akr1d1-/-*mice have increased insulin sensitivity and increased RER. Despite the 12α-hydroxylated/non-12α-hydroxylated ratio remaining low in *Akr1d1-/-*males on HFD, they were not protected against diet induced insulin resistance. It needs to be established whether specific bile acids differently regulate insulin sensitivity, or if the changes to bile acid composition is simply a biomarker reflecting a more complex mechanism.

Insulin sensitivity in *Akr1d1-/-*males was underpinned by increased protein expression of insulin signaling components in the liver (AKT) and skeletal muscle (AKT, INSRβ, mTOR). AKR1D1 knockdown also increased AKT1 gene and protein expression in Huh7 hepatoma cells, indicating autocrine regulation in the liver. Consistent with changes in bile acid composition contributing to the regulation of insulin signaling, AKR1D1 knockdown decreased CDCA levels to a greater degree than CA and the increase in *AKT1* expression was partially reversed by treatment with CDCA but not CA. CDCA has a greater affinity for FXR than CA^42^ and in our recent *in vitro* study AKR1D1 knockdown increased total and insulin stimulated AKT in HepG2 hepatoma cells and the FXR agonist GW4064 prevented the induction of AKT^19^. However, insulin tolerance tests are largely a measure of glucose uptake into muscle and as AKR1D1 is not expressed in myocytes systemic factors must be involved. Moreover, increased insulin stimulated glucose uptake was only seen in mature *Akr1d1-/-*mice, suggesting muscle insulin sensitivity develops over time, or that genotype interacts with age-related changes in bile acid^43^ or metabolic homeostasis^44^. Skeletal muscle expresses TGR5 (but not FXR) and transgenic mice that overexpress TGR5 in skeletal muscle have increased insulin sensitivity, secondary to increased glycolytic flux^45^. There is an increased relative abundance of potent TGR5 agonists (UDCA, CDCA and LCA) in the serum of male *Akr1d1-/-*mice and an increased RER making it possible that this underpins the improvement in insulin sensitivity.

In addition to insulin sensitivity, male *Akr1d1-/-*mice had a marked lipid phenotype, with reduced liver and adipose lipid accumulation concomitant with hypertriglyceridemia and high serum fatty acids. This was underpinned by downregulation of PPARγ and its transcriptional targets, suggesting reduced fatty acid uptake (*Cd36*), esterification (*Gpat3*) and storage (*Plin*4) in the liver. This phenotype is in contrast to *Cyp7a1-/-*mice which, although mirroring the insulin sensitivity of *Akr1d1-/-*males, have normal liver triacylglycerol and serum lipids^41^. In human hepatoma cells, the effects of AKR1D1 knockdown on the expression of lipid metabolism genes were partially recovered by LXRα, LXRβ and PXR antagonism as well as FXR agonism^18–20^ suggesting accumulation of oxysterols and bioactive intermediates of bile acid synthesis. However, in *Akr1d1-/-*males we did not observe any differences in 27-hydroxycholesterol or the AKR1D1 substrates 7α-12α-dihydroxy-4-chol-3-one and 7α-hydroxy-4-chol-3-one, and Ingenuity Pathway Analysis did not identify hepatic gene expression signatures associated with oxysterols, bile acids or bile acid intermediates. However, samples were taken in the fed state and the network of bile acid receptors and bile acid metabolizing enzymes are regulated in response to nutritional phase^46,47^ meaning studies investigating the transitions between feeding and fasting in the *Akr1d1-/-*mice is required. The predicted increase in STAT5B signaling is of interest as it is a master regulator of hepatic sexual dimorphism and negatively regulates both PPARγ and PPARα. AKR1D1 is also able to clear the sex steroids testosterone and androstenedione^17,48^, but whether this impacts androgen availability and androgen receptor (AR) activation is unknown. Circulating sex steroids were normal in *Akr1d1-/-*mice and pathway analysis did not predict AR activation. However, as there is cross-talk between AR, STAT5B^49^ and PPARγ^50^ signaling, whether increased androgen availability contributes to the metabolic effects of *Akr1d1* deletion warrants further investigation.

Whilst *Akr1d1-/-*males had a broad metabolic phenotype the effect on females was mild. In comparison to males the bile acid profile of *Akr1d1-/-*females was more similar to wildtype and the reduced 12α-hydroxylated/non-12α-hydroxylated ratio less pronounced. The expression and regulation of hepatic enzymes is highly sexually dimorphic^51,52^ and female specific changes in bile acid metabolism may help to limit the impact of *Akr1d1* deletion. Female *Akr1d1-/-*mice tended to have decreased 27-hydroxycholesterol (p=0.08) with significantly increased 7α-12α-dihydroxy-4-chol-3-one and 7α-hydroxy-4-chol-3-one, suggesting cholesterol is diverted from alternative, toward classic, synthesis. This female specific upregulation of classic synthesis is also observed in mice fed a high cholesterol diet or cholestyramine^53^. In the classic pathway, the observed upregulation of 12α-hydroxylase (CYP8B1) expression may help to maintain the production of CA. Bile acid detoxification pathways were also increased in females, and as sulfation and subsequent renal clearance of CDCA occurs at a rate twice that of CA^54,55^, enhanced CDCA clearance could also protect against the relative loss of 12α-hydroxylated bile acids. To date, no studies have investigated whether AKR1D1 plays a role in sex specific differences in metabolic homeostasis in humans.

The hepatic insulin sensitivity in *Akr1d1-/-*mice is consistent with that seen after AKR1D1 knockdown in human hepatoma cells^19^, however there is a discrepancy in the effect on lipid metabolism between the two models. In contrast to reduced hepatic triacylglycerol accumulation seen *in vivo*, AKR1D1 knockdown in human hepatoma cells increased triacylglycerol due to increased *de novo* lipogenesis and decreased β-oxidation^19^. The extent with which this reflects differences between species or between *in vitro* and *in vivo* systems is unclear. Bile acid profiles differ between mice and humans. In mice, CDCA is quickly converted to α or β-MCA and these primary bile acids, as well as their secondary metabolites, differ in their ability to activate or antagonize bile acid receptors. Although *in vitro* models are key in understanding autocrine regulation they cannot replicate secondary bile acid metabolism, sexual dimorphism, or tissue-tissue and endocrine interactions.

Bile acid metabolizing enzymes may have potential as therapeutic targets in liver disease. Although male *Akr1d1-/-*mice were lean with improved insulin sensitivity, they developed hypertriglyceridemia and were not protected against diet induced obesity or insulin resistance. AKR1D1 is downregulated in patients with type 2 diabetes^31^ and NAFLD^19^ and this may have implications beyond changes in energy metabolism. Bile acids and bioactive intermediates of their synthesis play important roles in proliferation, cytoprotection and immunoregulation^56,57^ and whether AKR1D1 plays in these processes needs to be further explored.

In conclusion, we have shown that AKR1D1 activity regulates insulin sensitivity and lipid metabolism *in vivo* and that its effects are sexually dimorphic. Further studies are clearly warranted to explore both the mechanisms by which this occurs and the role it plays in the pathogenesis of metabolic disease.

## Acknowledgments

This work was supported by the Medical Research Council, UK (programme grant awarded to J.W.T ref. MR/P011462/1); NIHR Oxford Biomedical Research Centre (Principal investigator award to J.W.T) based at Oxford University Hospitals NHS Trust and University of Oxford; Nigel Groome PhD Studentship awarded to L.L.G and A.A; Bioscientifica Trust Grant to N.N; Swiss National Science Foundation No 31003A-179400 (Principle Investigator A.O). The views expressed are those of the authors and not necessarily those of the NHS, the NIHR or the Department of Health.

## Supplementary Figures and Tables

**Supplementary Figure 1: Mature (30-week) *Akr1d1-/-*mice show no evidence of hepatic cholestasis, inflammation or damage**. Liver histology (H&E) showed no evidence of cholestasis (A) or hepatic inflammation (B). Serum levels of the markers of liver damage alanine aminotransferase (ALT) (C) and AST aspartate aminotransferase (D) were unchanged. Data are presented as mean±se of n=10-15 mice.

**Supplementary Figure 2: Intestinal lipid absorption and glucose control are apparently normal in young (10-week) *Akr1d1-/-*mice**. Fecal energy (A) and lipid content (B) are normal in *Akr1d1-/-*mice (n = 7 mice). Ip insulin tolerance (C), ip glucose tolerance (D), oral glucose tolerance (E), serum GLP-1 15 minutes post oral glucose bolus (F) and serum insulin 60 minutes post oral glucose bolus (G) were unchanged in male or female *Akr1d1-/-*mice (n = 15 mice). Data are presented as mean±se. *p<0.05 compared to wildtype of the same sex. (WT = wildtype C57BL/6; -/- = *Akr1d1-/-*)

**Supplementary Figure 3: Original Western blots from male quadricep muscle, liver and gonadal fat**. Original and complete Western blots used to quantify the protein expression of insulin signaling components insulin receptor β (INSRβ), insulin receptor substrate 1 (IRS1), protein kinase B (total-AKT) and mammalian target of rapamycin (mTOR) in male quadricep (A & B), liver (C & D) and gonadal fat (E & F). Data are presented as mean±se of n=3-10 mice. *p<0.05 compared to wildtype of the same sex. (WT = wildtype C57BL/6; -/- = *Akr1d1-/-*)

**Supplementary Figure 4: Original Western blots from female quadricep muscle, liver and gonadal fat**. Original and complete Western blots used to quantify the protein expression of insulin signaling components insulin receptor β (INSRβ), insulin receptor substrate 1 (IRS1), protein kinase B (total-AKT) and mammalian target of rapamycin (mTOR) in female quadricep (A & B) (n = 8 mice), liver (D & E) (n = 9-10 mice) and gonadal fat (G & H) (n = 6 mice). Quadricep and hepatic glycogen levels were unchanged (C & F) (n = 12 mice). Data are presented as mean±se. *p<0.05 compared to wildtype of the same sex. (WT = wildtype C57BL/6; -/- = *Akr1d1-/-*)

**Supplementary Figure 5: In female mice *Akr1d1* deletion reduces adipose mass but does not result in hypertriglyceridemia**. Female *Akr1d1-/-*mice (grey bars) have smaller gonadal and peri-renal adipose weights and increased intrascapular brown adipose depot weight compared to WT littermates (black bars) (A) (n = 8 mice). Serum HDL and total cholesterol (B), triacylglycerol (C), monoacylglycerols and diacylglycerols (D) are normal in Akr1d1-/-females but several fatty acid species are reduced (E) (n = 10 mice). Hepatic triacylglycerol (F) and glycogen (G) are unchanged. The expression of lipid metabolism genes in the gonadal fat (H) and liver (I) were unchanged (n = 10 mice). Data are presented as mean±se or log2(FC). *p<0.05 compared to wildtype. (WT = wildtype C57BL/6; -/- = *Akr1d1-/-*). mRNA expression was measured by RNASeq.

**Supplementary Table 1. Liver and serum bile acid concentrations in mature (30-week) wildtype and *Akr1d1-/-*mice on control diet**. Plasma bile acid concentration in liver and serum of wildtype and *Akr1d1-/-*mice. *p<0.05, * *p<0.01, * * *p<0.005, ^Ø^p<0.001, ^ØØ^p<0.0005, ^ØØØ^p<0.0001 compared to wildtype within sex. T, tauro; G, glycol; CA, cholic acid; CDCA, chenodeoxycholic acid; MCA, murocholic acid; DCA, deoxycholic acid; LCA, lithocholic acid; UDCA, ursodeoxycholic acid; HDCA, hyodeoxycholic acid; ND, not detected. Data are presented as mean±se of n= 11-15 mice.

**Supplementary Table 2. mRNA expression of bile acid metabolism, insulin signaling and lipid metabolism genes in the liver and quadricep muscle of mature (30-week) *Akr1d1-/-*mice**. Data are presented as mean±se of relative expression ratio, n=9-12 mice. *p<0.05 compared to wildtype. (WT = wildtype C57BL/6; -/-= *Akr1d1-/-*)

**Supplementary Table 3. Relative levels of serum lipids in (30-week) wildtype and Akr1d1-/-mice on control diet**. Metabolon metabolomics relative signal intensity for serum monoacylglycerol, diacylglycerol and fatty acids. *p<0.05, * *p<0.01, ^ØØ^p<0.0005, compared to wildtype within sex. Data are presented as mean±se of n=10 mice.

**Supplementary Table 4. Liver and serum bile acid concentrations in mature (30-week) wildtype and male *Akr1d1-/-*mice on high fat diet**. Plasma bile acid concentration in liver and serum of wildtype and *Akr1d1-/-*mice. *p<0.05, * *p<0.01, * * *p<0.005, ^Ø^p<0.001, ^ØØ^p<0.0005, ^ØØØ^p<0.0001 compared to wildtype within sex. T, tauro; G, glycol; CA, cholic acid; CDCA, chenodeoxycholic acid; MCA, murocholic acid; DCA, deoxycholic acid; LCA, lithocholic acid; UDCA, ursodeoxycholic acid; HDCA, hyodeoxycholic acid; ND, not detected. Data are presented as mean±se of n=11-15 mice.

