## Supplementary figures and images for "*Akr1d1-/-* mice have a sexually dimorphic metabolic phenotype with reduced fat mass, increased insulin sensitivity and hypertriglyceridemia in males"

### Supplemental Figures

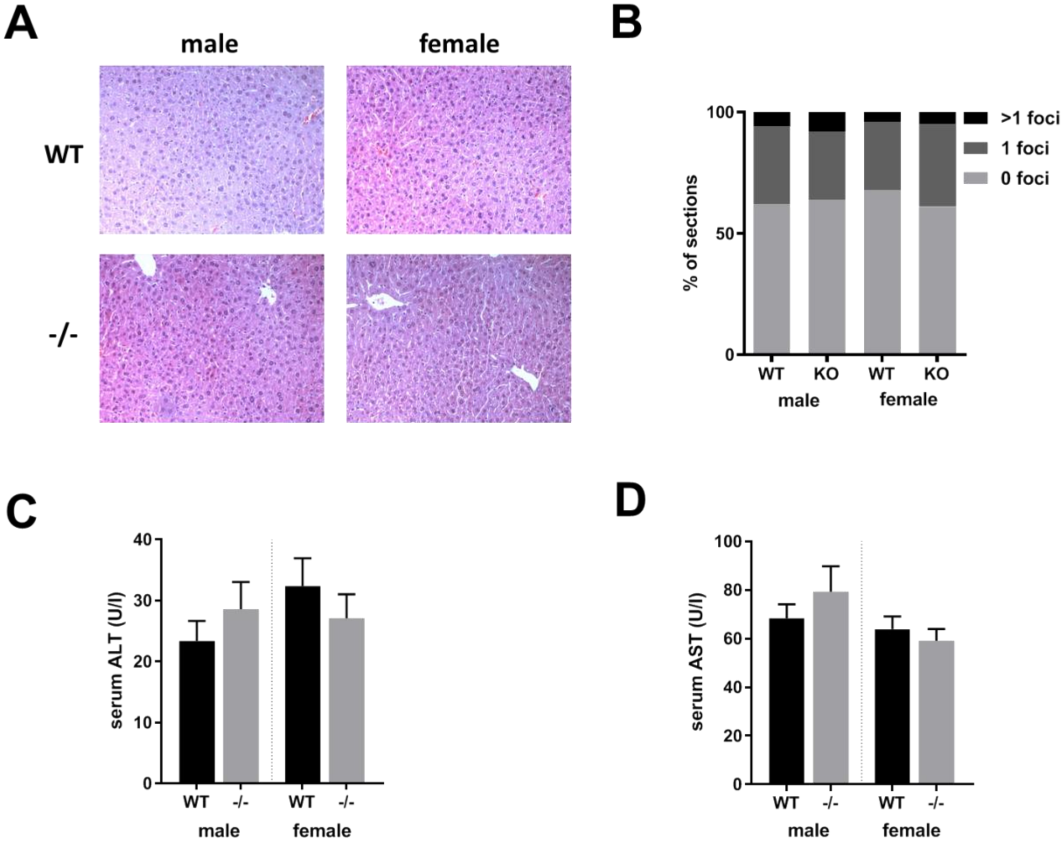

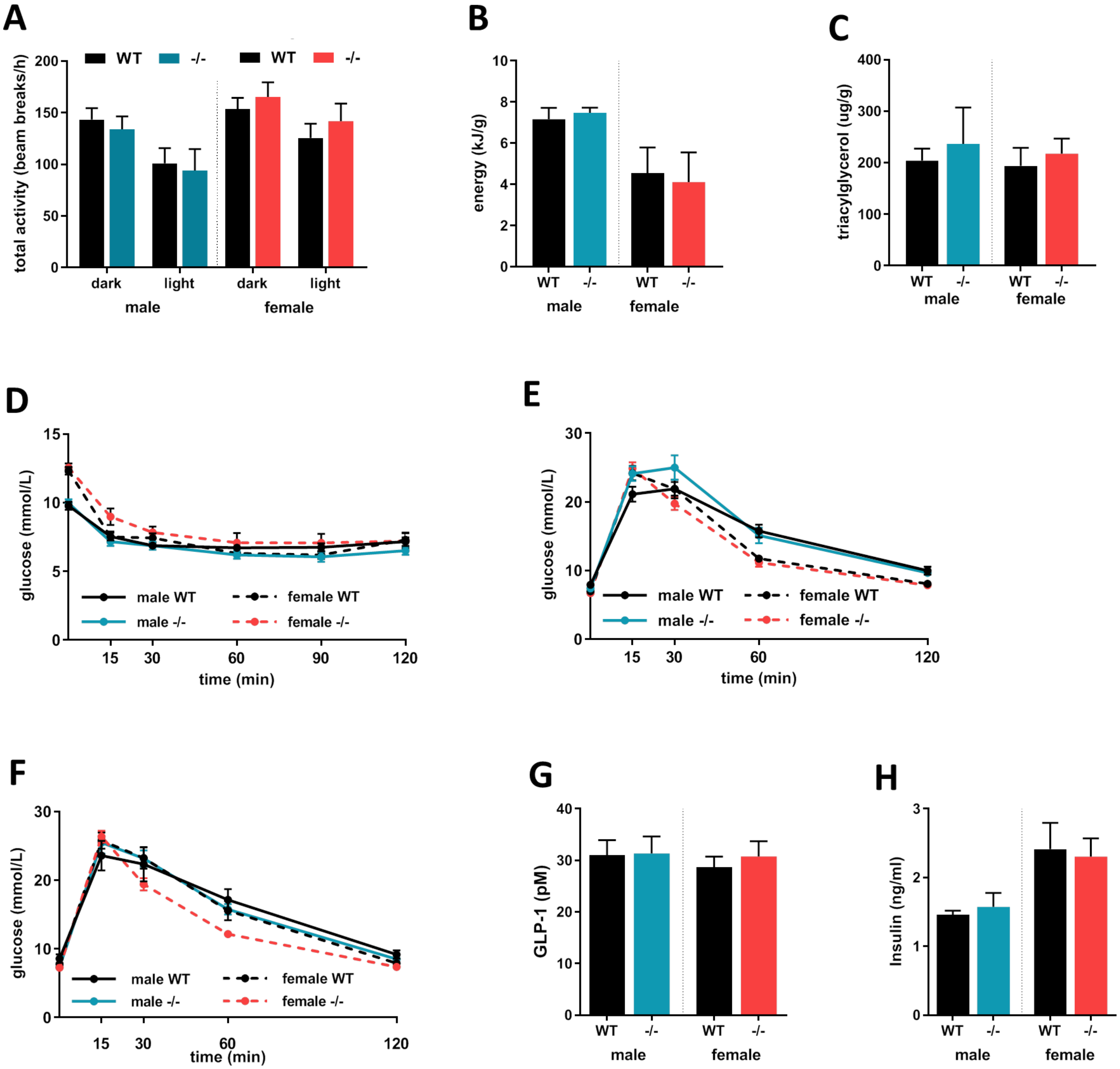

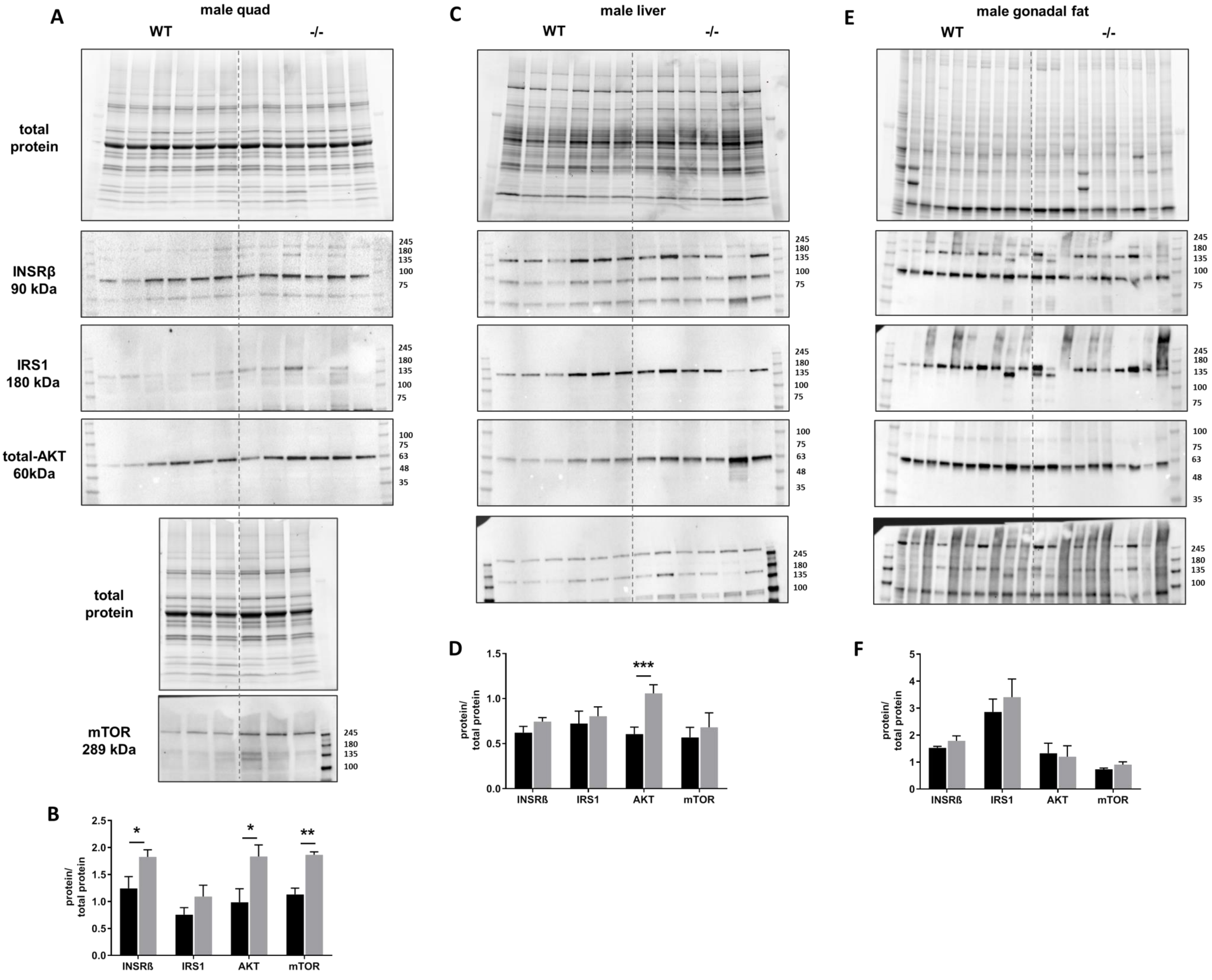

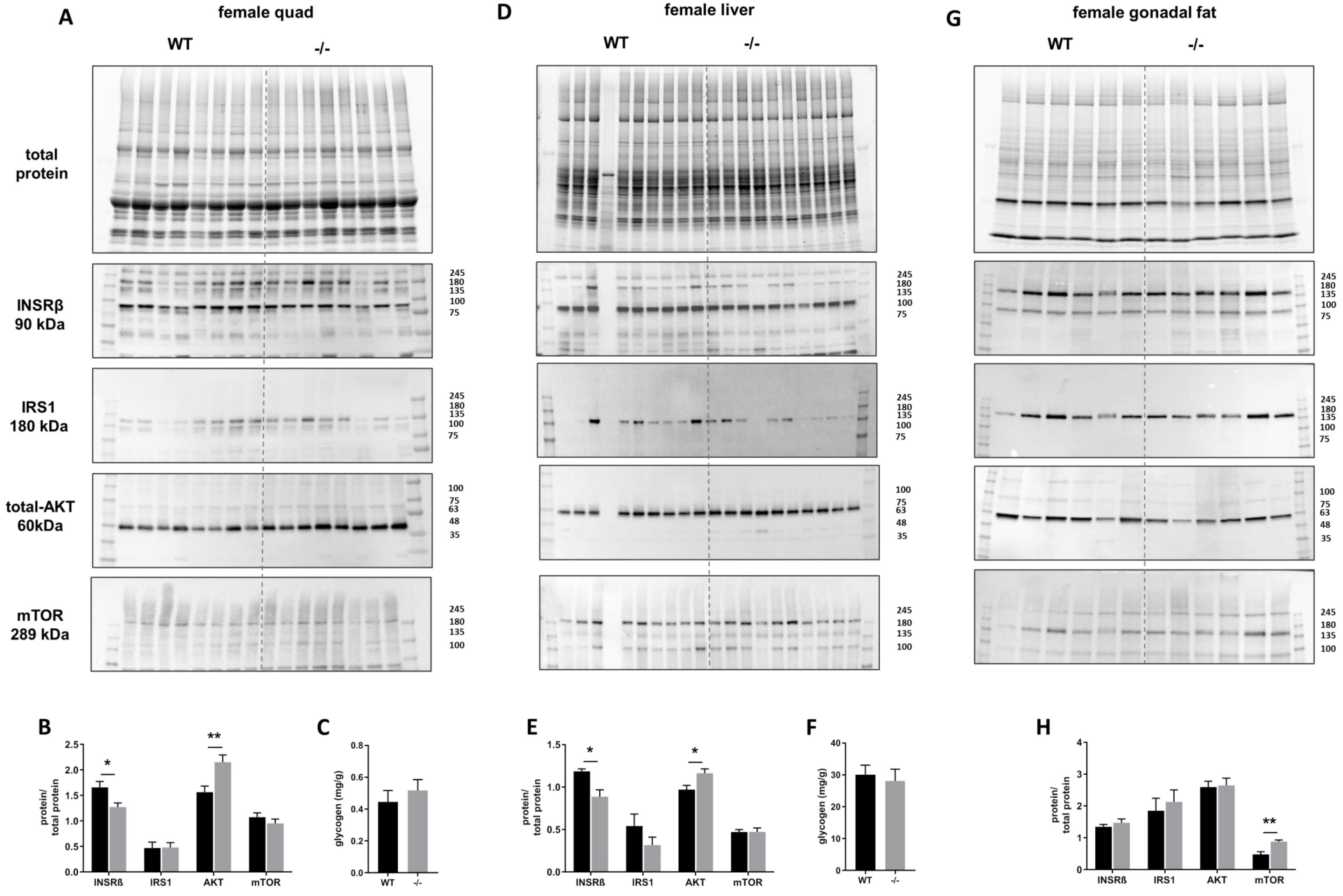

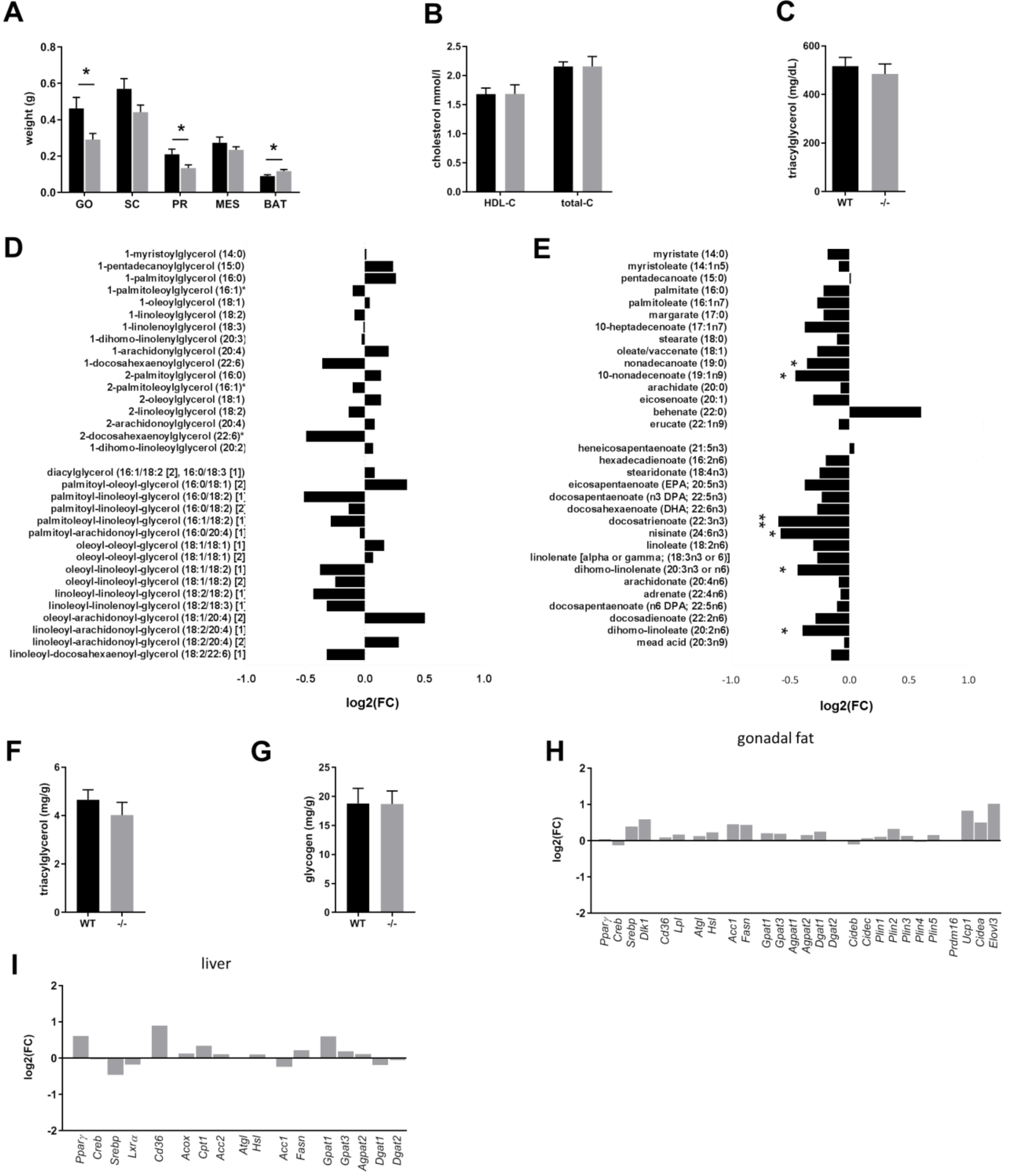
