## Supplemental Table 1 for "*Akr1d1-/-* mice have a sexually dimorphic metabolic phenotype with reduced fat mass, increased insulin sensitivity and hypertriglyceridemia in males"

| | liver (pmol/mg) | | | | serum (nM) | | | |
|---|---|---|---|---|---|---|---|---|
| | male | | female | | male | | female | |
| | WT | *Akr1d1-/-* | WT | *Akr1d1-/-* | WT | *Akr1d1-/-* | WT | *Akr1d1-/-* |
| **CA** | 5.23±1.88 | 0.12±0.06 | 6.26±2.58 | 1.22±0.53 | 365±112 | 49.1.6±20.4 | 837±177 | 246±90 |
| **T-CA** | 78.1±13.8 | 7.55±1.85$^{\phi}$ | 115±16.5 | 33.3±9.26*** | 92.3±16.3 | 5.1±5.1$^{\phi}$ | 417±47 | 143±24$^{\phi}$ |
| **G-CA** | 0.47±0.09 | 0.01±0.01*** | 0.47±0.08 | 0.21±0.03 | ND | ND | ND | ND |
| **CDCA** | 0.17±0.03 | ND$^{\phi}$ | 0.09±0.02 | ND | 31.2±4.8 | 19.2±2.3 | 33.3±4.3 | 18.7±1.9 |
| **T-CDCA** | 7.32±1.66 | 0.62±0.13** | 9.39±1.45 | 1.59±0.39$^{\phi}$ | 9.09±1.21 | 0.69±0.69$^{\phi\phi\phi}$ | 18.0±2.8 | 0.68±0.47$^{\phi\phi}$ |
| **G-CDCA** | ND | ND | ND | ND | 6.80±1.50 | 9.83±2.30 | 11.2±1.5 | 9.95±1.81 |
| **α-MCA** | 0.97±0.30 | 0.03±0.02* | 0.49±0.13 | 0.07±0.04 | 20.1±10.0 | 1.49±1.49 | 42.5±10.9 | 4.35±2.82* |
| **T-α-MCA** | 24.8±4.9 | 2.40±0.58*** | 27.6±4.0 | 4.89±1.32 | 46.7±10.4 | 7.9±1.7** | 107±13 | ND$^{\phi\phi\phi}$ |
| **β-MCA** | 4.87±1.20 | 0.85±0.24 | 1.62±0.35 | 0.54±0.14 | 111±39 | 24.1±8.1 | 450±114 | 147±47 |
| **T-β-MCA** | 74.9±13.3 | 18.4±3.9** | 97.0±11.3 | 42.7±8.2* | 56.4±13.9 | 11.36±3.7* | 169±15 | 98±37 |
| **ω-MCA** | 1.03±0.22 | 0.24±0.06 | 0.61±0.19 | 0.21±0.06 | 129±33 | 31.3±6.4 | 398±79 | 113±32* |
| **T-ω-MCA** | 60.3±12.4 | 20.7±4.5 | 67.2±9.97 | 32.8±6.9 | 190±27 | 62.2±11.8** | 560±38 | 246±40$^{\phi\phi}$ |
| **DCA** | 0.64±0.20 | ND | 0.28±0.03 | 0.02±0.02$^{\phi\phi\phi}$ | 443±77 | 37.4±15.4$^{\phi}$ | 1076±131 | 96.4±24.4$^{\phi\phi\phi}$ |
| **T-DCA** | 15.4±2.2 | 0.77±0.18$^{\phi\phi\phi}$ | 20.0±2.6 | 2.35±0.66$^{\phi\phi}$ | 58.7±9.9 | 1.83±1.5$^{\phi\phi\phi}$ | 156±22 | 3.39±2.02$^{\phi\phi\phi}$ |
| **LCA** | ND | ND | ND | ND | 77.5±3.7 | 68.7±5.8 | 65.8±3.1 | 65.5±4.8 |
| **T-LCA** | 1.47±0.07 | 0.99±0.24 | 1.85±0.13 | 1.94±0.22 | 0.65±0.16 | 1.21±1.04 | 0.69±0.27 | 1.35±0.38 |
| **G-LCA** | ND | ND | ND | ND | ND | ND | ND | ND |
| **UDCA** | 0.36±0.10 | 0.28±0.08 | 0.23±0.10 | 0.14±0.04 | 66.2±20.2 | 45.5±5.1 | 177±73 | 33.6±5.6 |
| **T-UDCA** | 10.1±2.1 | 7.49±1.23 | 12.8±2.01 | 7.77±1.94 | 42.6±3.75 | 20.0±3.4** | 94.2±7.6 | 61.0±11.0 |
| **G-UDCA** | 0.01±0.004 | ND | 0.01±0.004 | ND | 5.62±0.2 | 6.35±0.64 | 6.04±0.15 | 6.23±0.15 |
| **HDCA** | 0.62±0.01 | 2.28±0.44** | 0.08±0.03 | 0.18±0.05 | 54.9±18.2 | 28.4±5.1 | 125±21 | 5.38±2.13$^{\phi\phi\phi}$ |
| **7-oxo-DCA** | ND | ND | 0.09±0.05 | 0.03±0.02 | 20.1±13.8 | 40.0±11.3 | 25.6±11.4 | 140±101 |
| **T-7-oxo-LCA** | 0.10±0.04 | ND | 0.23±0.06 | 0.04±0.03 | ND | ND | ND | ND |
| **12α-hydroxyl** | 99.7±17.5 | 8.44±2.04 | 142±20 | 37.1±10.3 | 959±180 | 93±44 | 2486±299 | 489±113 |
| **non-12α-hydroxyl** | 187±34 | 55.3±10.4 | 219±27 | 92.9±18.5 | 868±178 | 379±44 | 2282±305 | 952±183 |
