## Supplemental Table 2 for "*Akr1d1-/-* mice have a sexually dimorphic metabolic phenotype with reduced fat mass, increased insulin sensitivity and hypertriglyceridemia in males"

| | male liver relative expression ratio | | female liver relative expression ratio | |
|---|---|---|---|---|
| | **WT** | ***Akr1d1-/-*** | **WT** | ***Akr1d1-/-*** |
| ***Cyp7a1*** | 0.37±0.08 | 0.25±0.08 | 1.62±0.41 | 2.51±0.80 |
| ***Cyp8b1*** | 0.93±0.10 | 0.69±0.05 | 0.60±0.15 | 1.34±0.21* |
| ***Cyp3a11*** | 0.59±0.11 | 0.63±0.22 | 1.61±0.20 | 2.18±0.28* |
| ***Sult2a7*** | 0.25±0.09 | 0.21±0.08 | 0.68±0.14 | 1.23±0.15* |
| ***Insrβ*** | 1.32±0.24 | 1.45±0.32 | 1.18±0.19 | 1.02±0.10 |
| ***Irs1*** | 1.14±0.08 | 1.07±0.20 | 1.21±0.32 | 1.33±0.26 |
| ***Pi3K*** | 0.48±0.06 | 0.50±0.10 | 0.22±0.03 | 0.30±0.32 |
| ***Akt1*** | 0.83±0.21 | 0.95±0.15 | 0.88±0.42 | 0.98±0.21 |
| ***mTor*** | 0.38±0.26 | 2.92±0.49 $^{\phi}$ | 0.62±0.08 | 0.75±0.17 |
| ***Pparγ*** | 0.92±0.13 | 0.60±0.07* | 0.32±0.35 | 0.39±0.34 |
| ***Cd36*** | 0.73±0.06 | 0.42±0.10* | 0.64±0.08 | 0.67±0.09 |
| ***Fasn*** | 0.47±0.05 | 0.31±0.15 | 0.55±0.09 | 0.43±0.13 |
| ***Gpat1*** | 0.84±0.17 | 0.32±0.43 | 0.73±0.14 | 0.84±0.32 |
| ***Gpat3*** | 0.79±0.14 | 0.32±0.09* | 0.61±0.14 | 0.66±0.17 |
| ***Cidec*** | 1.32±0.20 | 0.69±0.13* | 1.02±0.29 | 1.14±0.36 |
| ***Plin4*** | 0.63±0.09 | 0.24±0.16* | 0.59±0.25 | 0.48±0.10 |

| | male quad relative expression ratio | | female quad relative expression ratio | |
|---|---|---|---|---|
| | **WT** | ***Akr1d1-/-*** | **WT** | ***Akr1d1-/-*** |
| ***Insrβ*** | 1.18±0.07 | 2.51±0.80 | 1.13±0.07 | 1.00±0.05 |
| ***Irs1*** | 0.65±0.04 | 1.34±0.21* | 0.98±0.09 | 0.93±0.07 |
| ***Pi3K*** | 0.64±0.06 | 1.02±0.10 | 1.20±0.08 | 0.96±0.09$^{0.06}$ |
| ***Akt1*** | 1.15±0.08 | 1.33±0.26 | 1.14±0.07 | 0.98±0.06$^{0.09}$ |
| ***mTor*** | 0.43±0.10 | 0.30±0.32 | 0.37±0.08 | 0.53±0.13 |
| ***Cd36*** | 0.55±0.05 | 0.98±0.21 | 0.49±0.04 | 0.37±0.04$^{0.06}$ |
| ***Acc2*** | 1.09±0.08 | 0.75±0.17 | 1.10±0.07 | 0.96±0.10 |
| ***Cpt1*** | 1.07±0.06 | 0.39±0.34 | 1.16±0.10 | 1.07±0.07 |
| ***Acc1*** | 0.80±0.21 | 0.67±0.09 | 0.42±0.07 | 0.39±0.06 |
| ***Fasn*** | 0.99±0.25 | 0.43±0.13 | 0.37±0.08 | 0.53±0.13 |
| ***Dgat1*** | 1.34±0.22 | 0.84±0.32 | 0.80±0.13 | 0.86±0.09 |
| ***Gpam*** | 1.15±0.07 | 0.66±0.17 | 1.00±0.10 | 0.87±0.09 |
| ***Pdk4*** | 0.56±0.08 | 1.14±0.36 | 0.56±0.08 | 0.67±0.11 |
