## Supplemental Table 3 for "*Akr1d1-/-* mice have a sexually dimorphic metabolic phenotype with reduced fat mass, increased insulin sensitivity and hypertriglyceridemia in males"

| | male serum | | female serum | |
|---|---|---|---|---|
| | WT | *Akr1d1-/-* | WT | *Akr1d1-/-* |
| **monoacylglycerols** | | | | |
| 1-myristoylglycerol (14:0) | 0.86±0.057 | 1.1±0.071 | 1.1±0.11 | 1.1±0.24 |
| 1-pentadecanoylglycerol (15:0) | 1.1±0.099 | 1.2±0.081 | 1.0±0.095 | 1.2±0.26 |
| 1-palmitoylglycerol (16:0) | 0.98±0.12 | 1.2±0.093 | 1.0±0.13 | 1.3±0.29 |
| 1-palmitoleoylglycerol (16:1)* | 1.3±0.19 | 2.2±0.22* | 0.63±0.069 | 0.59±0.098 |
| 1-oleoylglycerol (18:1) | 1.0±0.16 | 1.4±0.13* | 0.75±0.061 | 0.78±0.082 |
| 1-linoleoylglycerol (18:2) | 1.0±0.092 | 1.6±0.19* | 0.94±0.078 | 0.88±0.16 |
| 1-linolenoylglycerol (18:3) | 0.99±0.098 | 1.4±0.14 | 0.98±0.11 | 0.97±0.23 |
| 1-dihomo-linolenylglycerol (20:3) | 1.2±0.24 | 1.6±0.14* | 0.76±0.071 | 0.75±0.10 |
| 1-arachidonylglycerol (20:4) | 1.3±0.17 | 1.4±0.14 | 0.91±0.079 | 1.0±0.15 |
| 1-docosahexaenoylglycerol (22:6) | 1.4±0.31 | 1.7±0.22 | 0.85±0.12 | 0.66±0.12 |
| 2-palmitoylglycerol (16:0) | 1.0±0.12 | 1.4±0.34 | 1.1±0.11 | 1.2±0.25 |
| 2-palmitoleoylglycerol (16:1)* | 1.3±0.18 | 2.0±0.20 | 0.61±0.068 | 0.57±0.092 |
| 2-oleoylglycerol (18:1) | 1.1±0.23 | 1.5±0.14* | 0.77±0.084 | 0.84±0.13 |
| 2-linoleoylglycerol (18:2) | 1.0±0.083 | 1.5±0.19* | 0.92±0.073 | 0.84±0.14 |
| 2-arachidonoylglycerol (20:4) | 1.0±0.17 | 1.2±0.24 | 0.71±0.13 | 0.76±0.085 |
| 2-docosahexaenoylglycerol (22:6)* | 1.4±0.33 | 1.4±0.27 | 0.8±0.16 | 0.76±0.085 |
| 1-dihomo-linoleoylglycerol (20:2) | 1.2±0.13 | 1.7±0.19 | 0.81±0.077 | 0.84±0.11 |
| **diacylglycerols** | | | | |
| diacylglycerol (16:1/18:2 [2], 16:0/18:3 [1]) | 1.1±0.29 | 1.6±0.28 | 0.33±0.67 | 0.35±0.12 |
| palmitoyl-oleoyl-glycerol (16:0/18:1) [2] | 0.80±0.19 | 1.4±0.22* | 0.28±0.033 | 0.35±0.092 |
| palmitoyl-linoleoyl-glycerol (16:0/18:2) [1] | 1.6±0.23 | 1.7±0.27 | 0.75±0.096 | 0.53±0.10 |
| palmitoyl-linoleoyl-glycerol (16:0/18:2) [2] | 1.1±0.22 | 1.7±0.32 | 0.62±0.096 | 0.56±0.18 |
| palmitoleoyl-linoleoyl-glycerol (16:1/18:2) [1] | 1.4±0.19 | 2.9±0.31$^{\phi\phi}$ | 0.59±0.064 | 0.49±0.071 |
| palmitoyl-arachidonoyl-glycerol (16:0/20:4) [1] | 1.1±0.24 | 1.1±0.22 | 0.56±0.074 | 0.54±0.13 |
| oleoyl-oleoyl-glycerol (18:1/18:1) [1] | 0.79±0.16 | 1.8±0.32** | 0.55±0.1 | 0.62±0.12 |
| oleoyl-oleoyl-glycerol (18:1/18:1) [2] | 1.2±0.22 | 2.2±0.37* | 0.7±0.13 | 0.73±0.11 |
| oleoyl-linoleoyl-glycerol (18:1/18:2) [1] | 1.2±0.16 | 1.7±0.15** | 0.78±0.066 | 0.6±0.087 |
| oleoyl-linoleoyl-glycerol (18:1/18:2) [2] | 1.0±0.14 | 1.9±0.16** | 0.77±0.058 | 0.64±0.12 |
| linoleoyl-linoleoyl-glycerol (18:2/18:2) [1] | 1.2±0.20 | 1.7±0.12 | 0.72±0.10 | 0.64±0.13 |
| linoleoyl-linolenoyl-glycerol (18:2/18:3) [1] | 0.77±0.19 | 0.72±0.13 | 0.87±0.10 | 0.58±0.11 |
| oleoyl-arachidonoyl-glycerol (18:1/20:4) [2] | 1.1±0.20 | 1.4±0.21 | 0.58±0.087 | 0.82±0.15 |
| linoleoyl-arachidonoyl-glycerol (18:2/20:4) [1] | 1.2±0.21 | 1.6±0.26 | 0.8±0.10 | 0.81±0.16 |
| linoleoyl-arachidonoyl-glycerol (18:2/20:4) [2] | 1.0±0.15 | 1.6±0.21* | 0.68±0.082 | 0.83±0.15 |
| linoleoyl-docosahexaenoyl-glycerol (18:2/22:6) [1] | 1.3±0.34 | 1.7±0.41 | 1.0±0.15 | 0.79±0.22 |
| **polyunsaturated fatty acids** | | | | |
| myristate (14:0) | 1.0±0.095 | 1.4±0.12* | 1.0±0.11 | 0.89±0.11 |
| myristoleate (14:1n5) | 1.1±0.22 | 1.5±0.21 | 0.89±0.15 | 0.83±0.14 |
| pentadecanoate (15:0) | 1.0±0.064 | 1.2±0.10 | 1.1±0.085 | 1.1±0.2 |
| palmitate (16:0) | 0.98±0.037 | 1.2±0.067* | 0.99±0.073 | 0.85±0.11 |
| palmitoleate (16:1n7) | 1.1±0.10 | 1.7±0.17* | 0.87±0.11 | 0.72±0.072 |
| margarate (17:0) | 0.99±0.037 | 1.1±0.067 | 1.1±0.099 | 0.96±0.14 |
| 10-heptadecenoate (17:1n7) | 1.0±0.043 | 1.4±0.089* | 1.1±0.13 | 0.81±0.059 |
| stearate (18:0) | 0.97±0.026 | 1.1±0.07 | 1.1±0.069 | 1.1±0.17 |
| oleate/vaccenate (18:1) | 0.95±0.076 | 1.3±0.095* | 1.1±0.11 | 0.95±0.08 |
| nonadecanoate (19:0) | 1.2±0.065 | 0.96±0.099 | 0.82±0.078 | 0.64±0.094* |
| 10-nonadecenoate (19:1n9) | 1.0±0.047 | 1.4±0.11* | 1.1±0.12 | 0.79±0.073* |
| arachidate (20:0) | 1.1±0.063 | 1.5±0.099* | 0.72±0.05 | 0.69±0.11 |
| eicosenoate (20:1) | 1.3±0.13 | 1.7±0.14* | 0.79±0.075 | 0.64±0.12 |
| behenate (22:0) | 1.9±0.57 | 2.1±0.51 | 1.2±0.33 | 1.9±0.55 |

| | | | | |
|---|---|---|---|---|
| erucate (22:1n9) | 1.6±0.22 | 2.4±0.22* | 0.5±0.05 | 0.48±0.13 |
| **polyunsaturated fatty acids** | | | | |
| heneicosapentaenoate (21:5n3) | 0.86±0.20 | 1.1±0.29 | 0.48±0.072 | 0.5±0.092 |
| hexadecadienoate (16:2n6) | 1.0±0.11 | 1.5±0.15* | 0.94±0.11 | 0.81±0.059 |
| stearidonate (18:4n3) | 0.85±0.07 | 2.0±0.37* | 1.0±0.092 | 0.84±0.068 |
| eicosapentaenoate (EPA; 20:5n3) | 0.98±0.096 | 1.4±0.15 | 1.1±0.096 | 0.83±0.087 |
| docosapentaenoate (n3 DPA; 22:5n3) | 1.2±0.14 | 1.7±0.23* | 0.88±0.068 | 0.75±0.066 |
| docosahexaenoate (DHA; 22:6n3) | 0.96±0.13 | 1.1±0.12 | 1.1±0.073 | 0.88±0.10 |
| docosatrienoate (22:3n3) | 1.4±0.092 | 2.4±0.25$^{\phi\phi}$ | 0.8±0.067 | 0.53±0.048** |
| nisinate (24:6n3) | 1.1±0.11 | 1.8±0.26** | 0.94±0.077 | 0.63±0.064* |
| linoleate (18:2n6) | 0.98±0.065 | 1.3±0.095 | 1.1±0.14 | 0.92±0.072 |
| linolenate [alpha or gamma; (18:3n3 or 6)] | 1.0±0.13 | 1.3±0.098 | 1.2±0.15 | 0.96±0.11 |
| dihomo-linolenate (20:3n3 or n6) | 1.1±0.099 | 1.3±0.12 | 1.1±0.098 | 0.79±0.053* |
| arachidonate (20:4n6) | 0.93±0.089 | 0.90±0.068 | 1.1±0.059 | 1.0±0.096 |
| adrenate (22:4n6) | 1.2±0.20 | 1.2±0.12 | 0.95±0.052 | 0.9±0.057 |
| docosapentaenoate (n6 DPA; 22:5n6) | 1.2±0.14 | 1.1±0.12 | 0.96±0.073 | 0.89±0.061 |
| docosadienoate (22:2n6) | 1.3±0.074 | 1.4±0.21$^{\phi\phi}$ | 0.64±0.053 | 0.87±0.071 |
| dihomo-linoleate (20:2n6) | 1.1±0.075 | 1.4±0.11* | 1.0±0.094 | 0.79±0.045* |
| mead acid (20:3n9) | 1.1±0.19 | 1.0±0.12 | 1.1±0.11 | 1.1±0.076 |
