## Supplemental Table 4 for "*Akr1d1-/-* mice have a sexually dimorphic metabolic phenotype with reduced fat mass, increased insulin sensitivity and hypertriglyceridemia in males"

| | liver (pmol/mg) | | serum (nM) | |
|---|---|---|---|---|
| | **WT** | ***Akr1d1-/-*** | **WT** | ***Akr1d1-/-*** |
| **CA** | 1.95±0.43 | 0.17±0.09** | 578±147 | 43.5±16.9 |
| **T-CA** | 80.3±8.5 | 17.2±10.6*** | 115±16.5 | ND |
| **G-CA** | 0.34±0.06 | 0.09±0.06 | ND | ND |
| **CDCA** | 0.14±0.02 | 0.01±0.01*** | 36.4±8.3 | 16.1±0.4 |
| **T-CDCA** | 6.78±0.53 | 1.43±0.73$^{\phi\phi\phi}$ | 35.6±9.5 | 0.49±0.47 |
| **G-CDCA** | ND | ND | 10.2±1.4 | 8.06±0.20 |
| **α-MCA** | 0.53±0.9 | 0.08±0.05*** | 25.3±9.81 | ND |
| **T-α-MCA** | 17.1±1.35 | 4.17±1.90$^{\phi\phi}$ | 172±52.4 | 11.0±4.15 |
| **β-MCA** | 2.56±0.36 | 0.78±0.23** | 76.0±28.0 | 12.5±5.7 |
| **T-β-MCA** | 38.3±3.9 | 13.8±4.3*** | 221±67 | 15.7±2.7 |
| **ω-MCA** | 0.56±0.10 | 0.13±0.03** | 94.5±34.9 | 9.82±3.38 |
| **T-ω-MCA** | 30.0±3.4 | 9.18±2.24$^{\phi}$ | 497±103 | 62.4±11.7 |
| **DCA** | 0.21±0.05 | ND** | 404±95 | 21.0±5.9 |
| **T-DCA** | 9.06±1.31 | 0.97±0.38$^{\phi\phi}$ | 76.3±13.4 | 2.95±1.44 |
| **LCA** | ND | ND | 79.3±8.9 | 72.2±5.5 |
| **T-LCA** | 1.18±0.07 | 0.53±0.14*** | 1.00±0.18 | ND |
| **G-LCA** | ND | ND | ND | ND |
| **UDCA** | 0.15±0.04 | 0.07±0.04 | 59.5±14.9 | 16.7±1.7 |
| **T-UDCA** | 4.97±0.58 | 2.69±0.62 | 71.3±13.4 | 17.0±3.1 |
| **G-UDCA** | ND | ND | 5.92±0.22 | 5.53±0.22 |
| **HDCA** | 0.34±0.06 | 0.58±0.14 | 39.7±8.3 | 16.1±0.4 |
| **7-oxo-DCA** | ND | ND | 18.2±13.4 | 26.0±12.4 |
| **T-7-oxo-LCA** | 0.04±0.01 | ND* | ND | ND |
| **12α-hydroxyl** | 91.8±10.4 | 18.5±11.1$^{\phi\phi}$ | 1815±434 | 67.5±24.0 |
| **non-12α-hydroxyl** | 103±10 | 33.4±10.4$^{\phi\phi}$ | 1443±299 | 298±59 |
